# Influenza A defective viral genomes and non-infectious particles are increased by host PI3K inhibition via anti-cancer drug alpelisib

**DOI:** 10.1101/2024.07.03.601932

**Authors:** Ilechukwu Agu, Ivy José, Abhineet Ram, Daniel Oberbauer, John Albeck, Samuel L. Díaz Muñoz

**Affiliations:** Department of Microbiology and Molecular Genetics, University of California, Davis, One Shields Ave, Davis CA 95616; Department of Molecular and Cellular Biology, University of California, Davis, One Shields Ave, Davis CA 95616; Genome Center, University of California, Davis One Shields Ave, Davis CA 95616

## Abstract

RNA viruses produce abundant defective viral genomes during replication, setting the stage for interactions between viral genomes that alter the course of pathogenesis. Harnessing these interactions to develop antivirals has become a recent goal of intense research focus. Despite decades of research, the mechanisms that regulate the production and interactions of Influenza A defective viral genomes are still unclear. The role of the host is essentially unexplored; specifically, it remains unknown whether host metabolism can influence the formation of defective viral genomes and the particles that house them. To address this question, we manipulated host cell anabolic signaling activity and monitored the production of defective viral genomes and particles by A/H1N1 and A/H3N2 strains, using a combination of single-cell immunofluorescence quantification, third-generation long-read sequencing, and the cluster-forming assay, a method we developed to titer defective and fully-infectious particles simultaneously. Here we show that alpelisib (Piqray), a highly selective inhibitor of mammalian Class 1a phosphoinositide-3 kinase (PI3K) receptors, significantly changed the proportion of defective particles and viral genomes (specifically deletion-containing viral genomes) in a strain-specific manner, under conditions that minimize multiple cycles of replication. Alpelisib pre-treatment of cells led to an increase in defective particles in the A/H3N2 strain, while the A/H1N1 strain showed a decrease in total viral particles. In the same infections, we found that defective viral genomes of polymerase and antigenic segments increased in the A/H1N1 strain, while the total particles decreased suggesting defective interference. We also found that the average deletion size in polymerase complex viral genomes increased in both the A/H3N2 and A/H1N1 strains. The A/H1N1 strain, additionally showed a dose-dependent increase in total number of defective viral genomes. In sum, we provide evidence that host cell metabolism can increase the production of defective viral genomes and particles at an early stage of infection, shifting the makeup of the infection and potential interactions among virions. Given that Influenza A defective viral genomes can inhibit pathogenesis, our study presents a new line of investigation into metabolic states associated with less severe flu infection and the potential induction of these states with metabolic drugs.

## Introduction

RNA virus infections frequently produce defective viral genomes, which can influence the course of infection through interactions such as complementation and interference (Dimmock, 2014). Non-clinical studies have extensively associated defective viral genome accumulation with reduced disease severity, while fewer but notable clinical studies have demonstrated this trend in Influenza A (Vasilijevic, 2017) and Respiratory Syncytial Virus (Felt, 2021) infections (Brennan, 2024). Consequently, defective viral genomes have become a recent focus of intense pre-clinical research (Smith, 2016; Meng, 2017; Wasik, 2018; Zhao, 2018; Bdier, 2019; Yamagata, 2019; Tapia, 2019; Harding, 2019), with their antiviral potential holding substantial implications for clinical applications and pandemic preparedness.

The study of the mechanisms that influence the production of defective viral genomes and the outcome of these virus-virus interactions during an infection have become a top priority to realize their public health potential. Influenza A virus infections are primarily composed of virions that cannot mount a complete infectious cycle. Only 1 – 30% of virions can propagate fully from cell to cell (Brooke, 2013; Brooke, 2017; Diefenbacher, 2018). While there are a variety of reasons why virions are not fully infectious—for example, SNPs or faulty protein expression—virions harboring genome segments with internal sequence deletions (Davis, 1980; Nayak, 1982; Saira, 2013) have been termed defective interfering particles (DIP) when their accumulation by *de novo* or exogenous means diminishes the productivity of Influenza A infections and leads to mild disease outcomes; a phenomenon termed *defective interference* (Dimmock, 2014; Manzoni, 2018; Vignuzi, 2019; Wu, 2022). These defective interfering particles do not produce the proteins necessary for a single DIP to complete an infection, leading to a non-propagative infection that relies on virus coinfection to disseminate (Yamagata, 2019). However, upon complementation, the viral genomes with internal deletions can actively interfere with the production of full-length viruses. Recently it has been shown that these “defective” viral segments can actually encode proteins (Ranum, 2024) that can also interfere with full length viral replication or that may have other functions. Thus, we will refer to these segments containing internal deletions as **Del**etion-containing **V**iral **G**enomes (hereafter DelVGs, after Alnaji et al. 2019). This rising relative abundance of DelVGs at the expense of full-length viral genomes (Dimmock, 2014; Ranum, 2024) has been associated with mild disease outcomes (Vasilijevic, 2017). The flood of complementation-dependent virions sets the stage for interactions that can alter the course of Influenza A pathogenesis, inspiring research into the factors that influence the generation of DelVGs.

Defective interfering particles and the genomes they harbor have been associated with high infection multiplicity (Von Magnus, 1954) and specific mutations of the viral polymerase (Rodriguez, 2013; Vasilijevic, 2017) or matrix genes (Perez-Cidoncha, 2014). At a molecular level, the best evidence on how DelVGs are generated (Alnaji, 2020) supports a model whereby the polymerase pauses synthesis while still processing template and resumes synthesis downstream, leading to an internal deletion (Nayak, 1982; Winter, 1981). The molecular determinants of this pause and why it occurs more frequently in some viral genome segments remains unknown (Alnaji, 2020). While it was dogma that a packaging advantage (Brooke, 2014; Alnaji, 2021; Meng, 2017; Odagiri, 1997) was the mechanism behind DelVG suppression of full-length virus (Brooke, 2014; Von Magnus, 1954; Pelz, 2021; Huang, 1970; Frensing, 2013; Alnaji, 2020), a recent study examining *de novo* intracellular DelVG generation in strain PR8 suggests that there is no packaging advantage (Alnaji, 2021). Studies have shown that both the DelVG RNA and its encoded proteins contribute to the suppression of full-length virus (Meng, 2017; Dadonaite, 2019; Hara, 2013; Octaviani 2011, Ranum, 2024). Notably, a recent study suggests that protein production from DelVGs was considerable and that some of these proteins competed for binding with their full-length cognates, impairing polymerase function (Ranum, 2024). Thus, considerable effort has gone into investigating the viral factors that influence DelVG production and the effects of DelVGs on the host (Wang, 2023). However, an almost completely unexplored potential modulator of DelVG production is the host cell.

Host cell variables, particularly metabolism and metabolic signaling, can affect progeny virus yield and the severity of infection. A comprehensive review of the impact of extreme nutritional states—such as caloric restriction and diet-induced obesity—on flu infection found eroded immune response and survivability in mice (Gardner, 2011). In humans, non-obese patient groups routinely shed lower progeny viral loads relative to obese patient groups (Honce, 2019; Honce, 2020). At a cellular level, reprogramming host tricarboxylic acid cycle with excess malonate diminishes total viral progeny yield in a dose-dependent manner (Ackermann, 1951). Influenza A itself has adaptations to steer host metabolism in a manner that facilitates productive infection, further proof of the outsized role of host metabolism in influenza pathogenesis. Specifically, some flu strains have specific mutations that affect PI3K, a crucial upstream gatekeeper of pro-growth signal transduction networks (Wee, 2017; Hopkins, 2020). A highly-conserved Y89 residue on the influenza NS1 effector protein has a selective and inhibitory interaction with the SH2 domain of host p85β—the regulatory subunit of Class 1a phosphoinositide-3-kinase (PI3K)—which unleashes the catalytic p110α subunit to aberrantly activate PI3K signaling in the absence of a bona fide signaling growth factor like insulin (Hale, 2006; Li, 2008). Thus, the action of NS1 contorts host metabolism into a state characterized by increased pools of the very precursor metabolites (Luo, 2018; Saha, 2014; Al-Saffer, 2010) necessary for uninterrupted biosynthesis of virion components, facilitating pathogenesis (Hale, 2006; Li, 2008). Inhibiting NS1::p85β with ΔNS1(Y89F) expectedly diminishes viral-induced PI3K activation and progeny yield (Hale, 2006). Because diminished viral yield can result from DelVG accumulation and the inducers of DelVG production remain unknown, it is a reasonable secondary hypothesis that NS1::p85β inhibition—or other form of PI3K inactivation—diminishes viral yield wholly or in part via the induction of DelVG production. However, to our knowledge, the impact of host metabolic signaling on the production of Influenza A defective particles and DelVGs has not been studied.

Given the well-established crosstalk between host metabolism and Influenza A pathogenesis, it is surprising more research has not focused on the role of hosts in shaping DelVG production. A key barrier to the pursuit of this missed opportunity is the lack of tools that can quantitatively measure the virion and genome composition of infections. To fill this gap, we implemented two tools. First, we developed the cluster-forming assay to simultaneously titer defective and fully infectious Influenza A virions by modifying the well-known immunofocus assay (Baker, 2013) and implementing a computational pipeline that automated analysis of fluorescent microscopy images. Second, to quantify DelVGs and full-length genomic segments simultaneously, we used long-read sequencing of whole genome amplicons with unique molecular identifiers (UMI) to enable read de-duplication (Routh, 2013; Jaworski, 2017; Sotcheff, 2023; Karst, 2021). Armed with these tools, knowledge of flu-induced PI3K-AKT pathway activation, and single-cell phospho-AKT (pAKT) measurement as a readout for PI3K network activity, we explored if alpelisib—a highly selective small molecule inhibitor of PI3K (Furet, 2013; Fritsch, 2014; Yang, 2019)—affected defective virion and DelVG production. We hypothesized that disrupting flu-mediated activation of PI3K signaling with alpelisib would increase defective virion and DelVG production.

We found that alpelisib treatment affects DelVG production in both circulating human Influenza A virus subtypes and changes the infectivity of virions generated during infections. We first confirmed that alpelisib suppresses PI3K network signaling activity in MDCK-London cells and that this treatment overrides virus-induced PI3K network signaling upregulation during infections. Under these conditions, alpelisib increased DelVG production in polymerase complex segments in the A/H1N1 strain and increased the average deletion size in A/H1N1 and A/H3N2 strains. Moreover, there was compelling evidence for defective interference in A/H1N1 infections as DelVGs increased while total viral genomes decreased. At the virion level, the proportion of non-infectious particles produced by the A/H3N2 strain infections was significantly increased, while the total number of viral particles in the A/H1N1 strain was significantly altered. Collectively, these results suggest that host Class 1a PI3K metabolic signaling receptor inactivation affects the outcome of Influenza A virus infections, steering the population towards more non-infectious particles and increased DelVG production. Our results highlight the importance of host cell factors in determining the outcome of influenza virus infections, potentially informing host metabolic states that predict infection outcomes (Engels, 2017), as well as therapeutics that can induce host-mediated changes towards mild infection outcomes.

## Methods

### Cells and Viruses

We obtained MDCK-London cells from the Unites States Centers for Disease Control and Prevention (CDC) Influenza Reagent Resource (IRR). We maintained cells in minimum essential media (MEM) plus 5% fetal bovine serum (FBS). Egg-passaged wildtype A/California/07/2009(H1N1) and A/Texas/50/2012(H3N2) influenza strains were a gift from the lab of Dr. Ted Ross. These initial stocks were double plaque purified in MDCK cells (ATCC/BEI) and propagated thereafter at low multiplicity of infection (MOI = 0.001) in MDCK cells (ATCC/BEI).

### Alpelisib Dosing Assay

To confirm that alpelisib inhibits PI3K network signaling in MDCK-London cells, we seeded MDCK-London cells overnight at low density in MEM plus 5% FBS media for 24 hr into collagen-treated, glass-bottom 96-well tissue culture plates. We then serum-weaned the partially confluent monolayers for 24 hr in MEM plus 5% bovine serum albumin (BSA). Three hours prior to conclusion of serum weaning, we spiked a 10 uL pre-treatment of DMSO vehicle control or 21X alpelisib directly into 200 uL of the serum weaning supernatant to reach a 1X concentration. At the end of serum weaning, we mock-infected monolayers with MEM plus 2% BSA and 1% Anti-Anti (Virus Infection Media; VIM) or virus-infected at a multiplicity (MOI) of 1 in VIM; no trypsin was used. As part of the inoculation regimen, we spiked 1.9 uL of DMSO or 21X alpelisib into 40 uL of the inoculum supernatant for a 1X concentration, in order to sustain drug effects throughout the virus-monolayer adsorption period. After the 1 hr adsorption incubation, we aspirated inocula, washed monolayers and topped with VIM, and we spiked 10 uL of DMSO or 21X alpelisib directly into 200 uL of VIM supernatant for a 1X concentration. After 17 hr p.i., we harvested and titrated supernatants to determine fully infectious (i.e., propagation-capable) and defective (i.e. propagation-incapable) progeny virus yield via the novel cluster-forming assay (see Methods and Supplementary Materials). We fixed monolayers, conducted immunofluorescence (IF) staining, and imaged to derive cellular-level phospho-AKT signal intensity (see Methods) as a readout for PI3K network signaling activity. We ran three biological replicates of the experiment on different days.

### Single-cell Dose Response pAKT Immunofluorescence Quantification

At the end of the alpelisib-treated flu infections, we fixed monolayers with 4% paraformaldehyde (PFA). Primary staining was carried out with rabbit monoclonal antibodies targeting pAKT(S473) (Cell Signaling mAb#4060) and mouse monoclonal antibodies targeting Influenza A nucleoprotein (Millipore Sigma MAB8257). Secondary staining was respectively carried out with fluorophore-conjugated goat-anti-rabbit (Thermo Fisher Scientific A-21245) and goat-anti-mouse (Jackson ImmunoResearch 115-645-062) antibodies. We imaged IF-stained monolayers on a Andor Zyla 5.5 scMOS camera and a 20x/0.75 NA objective microscope. Images were then processed to derive cellular-level pAKT, and nucleoprotein signal intensities. The image data were stored as .nd2 files and retrieved using the Bio-Formats toolbox for MATLAB, which can be obtained from www.openmicroscopy.org/bio-formats. Subsequently, a specialized MATLAB cell segmentation pipeline (Pargett, 2017) was employed to process the images. Briefly, this pipeline utilized Hoechst 33342 as nuclear markers to identify the nuclei of individual cells. After cell identification and segmentation, single-cell pAKT signal was quantified by calculating the average pixel intensities within each individual cell. These intensity values were then background subtracted. To measure the background signal intensity, a well without any cells was imaged. The MATLAB pipeline output florescence for each cells in AU units. To control for any biases in image selection, we randomly subsampled a third of the data set prior to analyses reported; results were comparable with the full data set (see code for details https://github.com/pomoxis/Alpelisib-SIP).

### Cluster-forming Assay: Titration of Fully Infectious and Defective Virions

To determine the titer of fully infectious (i.e., propagation-capable) and defective (i.e. propagation-incapable) virions simultaneously, we developed the cluster-forming assay. This assay combines aspects of the conventional plaque assay with the immunocytochemical staining and microscopy, as done by Brooke et al (2013). Specifically, the cluster-forming assay employs a low-viscosity overlay medium that remains in a semi-solid state, restricting diffusion of progeny virus to directly adjacent cells, much like a plaque assay. This low viscosity overlay is removable, so that monolayers can be fixed, stained with IF antibodies, and imaged like an immunofocus assay. The basic principle is that virions that were fully infectious would spread from cell to cell, forming clusters of florescence, while virions that were unable to spread would appear as individual foci (Brooke et al., 2013). We build on previous similar assays (Brooke et al. 2013) by automating the image processing and analysis to identify productive and abortive infections.

We briefly describe this assay below and provide detailed methods in Supplement: We seeded MDCK-London cells overnight (24 hr) at high density in MEM plus 5% FBS media into collagen-treated, glass-bottom 96-well tissue culture plates. We inoculated the confluent monolayers with serial dilutions of virus-borne supernatant and incubated for 1 hr to facilitate virus-monolayer adsorption, after which we aspirated the inoculum, washed monolayers with VIM, and overlaid monolayers with medium-viscosity culture medium (VIM plus 4% carboxymethyl cellulose and 1 ug/mL TPCK-Trypsin). At 11 h.p.i. we aspirated overlay medium and fixed monolayers with 4% PFA. We stained fixed monolayers with fluorophore-conjugated ICC/IF antibodies targeting Influenza A nucleoprotein (NP) and counter-stained with Hoechst. We imaged IF-stained monolayers on a fluorescence microscope to reveal the number of productive (i.e. infections that spread from cell-to cell) and non-productive units (i.e., infections that did not spread from cell-cell) infection events, respectively depicted by a cluster of infected cells (productive clustering unit/PCU), or solitary infected cells (non-clustering unit/NCU) (**Figure 4, Figure S2.1-2.5**).

To quantify PCUs and NCUs, we utilized MATLAB’s image processing toolbox. Immunofluorescence images were processed to create object masks for each unit, and nuclear segmentation was performed via the Hoechst signal (**Figure S2.6-2.7**). Masks were refined and filtered, and the number of cells within each unit was determined (**Figure S2.8**). The R Programming Language was employed to classify clusters as PCUs or NCUs based on size, followed by calculation of NCU titer and proportion.

### Viral Genome Sequencing by Nanopore Long-read Sequencing

To derive the genomic sequences of progeny Influenza A virus from our treatments, we began by isolating viral genomic RNA from 100 uL of treatment group supernatants (Zymo Research, Quick-DNA/RNA Viral MagBead kit R2140). Next, we used a 2-cycle RT-PCR reaction (Invitrogen Super-Script™ III One-Step RT-PCR System with Platinum™ Taq DNA Polymerase kit 12574026) to reverse transcribe viral genomic RNA into the first cDNA strand (1st PCR cycle), and then synthesize the second cDNA strand (2nd PCR cycle). The RT reaction to produce the first cDNA strand was primed with a 45bp forward primer (Integrated DNA Technologies) that included a complementary sequence to the uni12 region shared by all flu genomic segments (12bp), flanked with a unique molecular identifier (UMI) sequence (12bp) and a landing pad sequence for downstream barcoding primers (21bp): fwd 5’-TTTCTGTTGGTGCTGATATTGNNNNNNNNNNNNAGCRAAAGCAGG-3’. The PCR reaction to generate the second cDNA strand was primed with a 47bp reverse primer that included a complementary sequence to the uni13 region shared by all flu genomic segments (13bp), flanked by a UMI sequence (12bp) and the barcoding primer landing pad sequence (22bp): rev 5’-ACTTGCCTGTCGCTC-TATCTTCNNNNNNNNNNNNAGTAGAAACAAGG-3’. We used AMPure XP beads (Beckman Coulter, AMPure XP A63881) with manufacturer’s instructions to remove excess primers, followed by a 17-cycle amplification PCR (Invitrogen, Platinum SuperFi Master Mix 12358-050) of the umi-tagged reads with barcoding primers (Oxford Nanopore Technologies, PCR Barcoding Expansion 1-96 kit EXP-PBC096). The low number of cycles was designed to minimize PCR duplicates. We then pooled 60 ng of barcoded amplicons from each sample, cleaned and concentrated this pooled sample (Zymo Research, Select-A-Size DNA Clean & Concentrator D4080), and prepared a sequencing library in accordance with manufacturer instructions (Oxford Nanopore Technologies, Ligation-Sequencing-Kit-V14 SQK-LSK114). We loaded the pooled libraries into an R10 flow cell connected to a MinION MkIB device and ran a 72 hr sequencing protocol from the MinKNOW control software. Upon sequencing run termination, we used the Guppy basecaller software to barcode-demultiplex sequenced reads into their respective treatment groups.

### Classification and Quantification of DelVGs and Full-length Viral Segments

We describe the pipeline in brief below; the code for the pipeline and analyses is posted on GitHub (https://github.com/pomoxis/alpelisib-SIP). Demultiplexed amplicon sequences underwent quality control pre-processing prior to deduplication into representative sequences, after which representative sequences were classified into subgroups for DelVGs and full-length, standard viral genomes (SVG).

#### Quality Control

Our sequencing library preparation strategy began with a 1-cycle each RT-PCR then PCR addition of 12bp-long UMI sequences to the 3’ and 5’ termini of viral genomic RNA, followed by PCR addition of sequencing barcodes to both termini:

5’-barcode—spacer—landing.pad—UMI—uni12—*locus*—uni13—UMI—landing.pad—spacer—bar-code-3’

For quality control, we trimmed off barcode and barcode landing pad regions with C*utadapt*, then used *Cutadapt* once more to filter-in only amplicons with a 12bp-long UMI region. Finally, we confirmed the presence of well-formed uni primer regions in the filtered amplicons before advancing to UMI deduplication.

#### UMI Deduplication

We used *UMI-Tools* to sequentially group PCR duplicates by UMI, and then collapse them into a single representative read. In the final quality control step, we trimmed the uni primer region off representative reads with *Cutadapt*. By integrating UMI-deduplication into our workflow, we’ve mitigated the impact of PCR amplification bias on sequencing depths. Consequently, our delVG and SVG count data represent a quantitative measurement of the abundance of RNA molecules (genome segments) from which the amplicons were derived.

#### DelVG Characterization

UMI-deduplicated fastq files containing read sequences were processed with the Virus Recombination Mapping (*ViReMa*) software to identify recombination events per genomic read using the following parameters:

*--Seed 25 --MicroInDel_Length 20 --Aligner bwa --ErrorDensity 1,25*

Additionally, the *-ReadNamesEntry* switch was included in a separate ViReMa run of the same dataset in order to assign read name information to each recombination, which allowed us to collapse deletion events with the same read name into a single delVG observation with in-house Bash and AWK scripts:

*--Seed 25 --MicroInDel_Length 20 --Aligner bwa --ErrorDensity 1,25 -ReadNamesEntry*

#### Full-length Viral Genome Characterization

To characterize SVGs, we began by using the *bwa* alignment tool to determine the properties of reads and their alignment to the reference genome; this information is captured in the bitwise FLAG field (column 2) of the output SAM file. Next we used the *AWK* program to select only reads with proper alignment to the forward and reverse strands of reference genome—bitwise FLAGs 0×0 (0) and 0×10 (16) respectively—and used *AWK* yet again to filter reads that were within ±100bp the length of the reference genomic segment.

### Statistics

In general, we relied on linear or linear mixed models to test significance between treatments using base R and the nmle packages, respectively. Owing to the intrinsic heterogeneity of flu infections (Russell, 2018; Wang, 2020), we included bioreplicates as a random factor, unless otherwise indicated. We tested alpelisib inhibition of PI3K network signaling and its dose dependence, using a linear model and Dunnett’s contrasts. We tested influenza activation of PI3K network signaling activity using ANOVA and Tukey’s HSD, to test for differences among strains and the mock infection. We tested whether alpelisib affects the production of defective virions, by testing for differences in the proportion of non-clustering units (NCU) according to each alpelisib dose using a linear mixed model. We similarly tested for differences in the total viral particles detected by the cluster-forming assay. Finally, we tested whether alpelisib increases defective viral genome production by examining the proportion of DelVGs in each alpelisib concentration using a linear mixed model that controlled for viral genome segment identity, as these have documented differences in the production of DelVGs during infection. To determine if there were differences in the type of DelVGs generated during infections with alpelisib, we examined the size distribution of unique deletions (DelVGS species) using linear mixed models for each viral genome segment (which have known differences in DelVG production) compared to the mock. Code for statistics and analyses is posted on GitHub (https://github.com/pomoxis/alpelisib-SIP).

## Results

### 1. Alpelisib inhibits PI3K network signaling in MDCK-London cells

We set out to confirm that alpelisib inhibits PI3K network signaling in the MDCK-London cell line, as has been demonstrated in numerous other mammalian cell lines and cell-free assays (Furet, 2013; Fritsch, 2014; Yang, 2019). We leveraged immunocytochemistry and fluorescence microscopy to quantify pAKT activity at the single cell level (Pargett, 2017). We found strong evidence that alpelisib inhibited PI3K network signaling by measuring the activity of the downstream pAKT effector. Increased doses of alpelisib, across a broad 1.25 - 40 µM concentration range, resulted in a clear dose dependent decrease in pAKT activity (Adjusted R^2^ = 0.1672, p < 0.00001, **Figure 1A**), with a 4.5342 decrease in AU per µM of alpelisib. In qualitative terms, pAKT activity was clearly downregulated in the alpelisib-treated monolayer (**Figure 1B**) relative to the vehicle-treated monolayer (**Figure 1C**); whose baseline pAKT activity was dusted across the cytoplasm of some cells and focused in the nuclei of others. Expectedly, the insulin-treated positive control group showed vivid pAKT upregulation, wherein pAKT activity localization went past cytoplasm and nuclei to include the plasma membrane (**Figure 1D**, **Supplementary Figure S1**).

**Figure 1.**
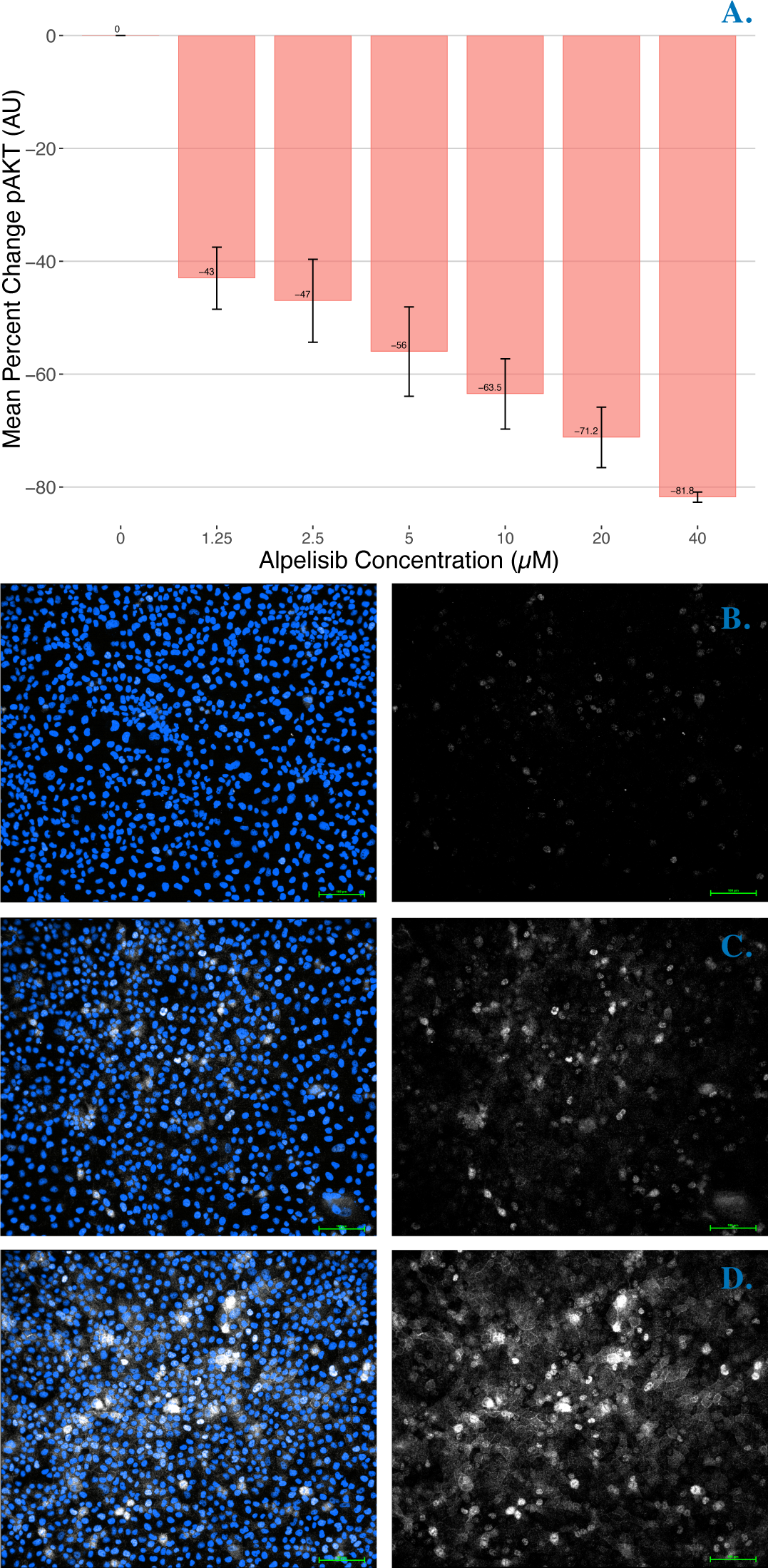
Alpelisib decreases pAKT activity (AU) of MDCK-London in a dose dependent manner. (**A**) Mean percent change in pAKT activity (AU) of MDCK-London cells exposed to increasing concentrations of alpelisib; 0 µM alpelisib treatment group received vehicle solvent (DMSO). n = 3 bioreplicates, sem. Fluorescence microscopy images of MDCK monolayers (**B**-**D**). (**B**) 20 µM alpelisib treated monolayer, (**C**) Vehicle treated monolayer, (**D**) Insulin treated monolayer (positive control). White/Cy5 – pAKT(S473); Blue/Hoechst – MDCK-London nucleus; green scale bar = 100µM.

### 2. Influenza A infection activates PI3K network signaling activity with strain-specificity in MDCK Cells

Class 1a PI3K signaling in host cells can be activated by several Influenza A strains, including A/WSN/1933(H1N1), A/Udorn/72(H3N2), A/Victoria (Hale, 2006; Ayllon, 2012), A/Puerto Rico/8/34(H1N1) (Hale, 2006; Li, 2008; Lopes, 2017), and an unspecified 1918 pandemic H1N1 strain (Cho, 2020). The effector domain of the viral NS1 effector protein is the molecular activator of the PI3K signaling cascade (Hale, 2006; Ayllon, 2012; Cho, 2020), and different influenza strains have been shown to differentially activate PI3K signaling in a tumorigenic cell line (Ayllon, 2012). To avoid confounding our results with the aberrant PI3K signaling typical in cancer cells (Fruman, 2014; Yuan, 2008; Fruman, 2017; Jokinen, 2015), and considering the strain-specific differences in the NS1 effector domain (**Figure 2**), we needed to ensure our chosen strains could activate PI3K network signaling in a non-tumorigenic MDCK cell line within the experiment window.

**Figure 2.**
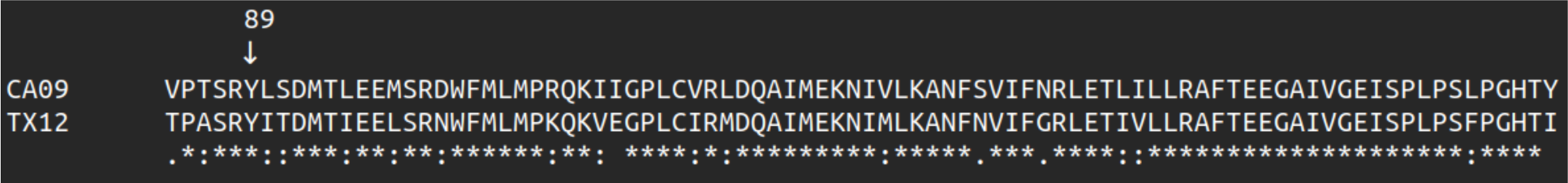
CA09 and TX12 have the highly conserved Y89 residue that is necessary for Class 1a PI3K activation. (**A**) Multiple sequence alignment of CA09 and TX12 NS genome segment sequences.

We measured pAKT in CA09 and TX12 infections compared to a mock infection. Both viral infections upregulated pAKT compared to mock infection in a statistically significant way (ANOVA Adjusted R^2^ = 0.2428, p < 0.00001), with CA09 and TX12 increasing the AU by an average of 205.861 and 106.886 respectively (Tukey Contrasts p < 0.00001). TX12 upregulation of pAKT was on average −98.976 AU less than CA09 and this effect was statistically significant (Tukey Contrasts p < 0.00001) (**Figure 3**). In addition to confirming PI3K activation by CA09, we discovered this trait in TX12 and demonstrated the differential dysregulation of PI3K signaling by both strains in a non-tumorigenic cell line.

**Figure 3.**
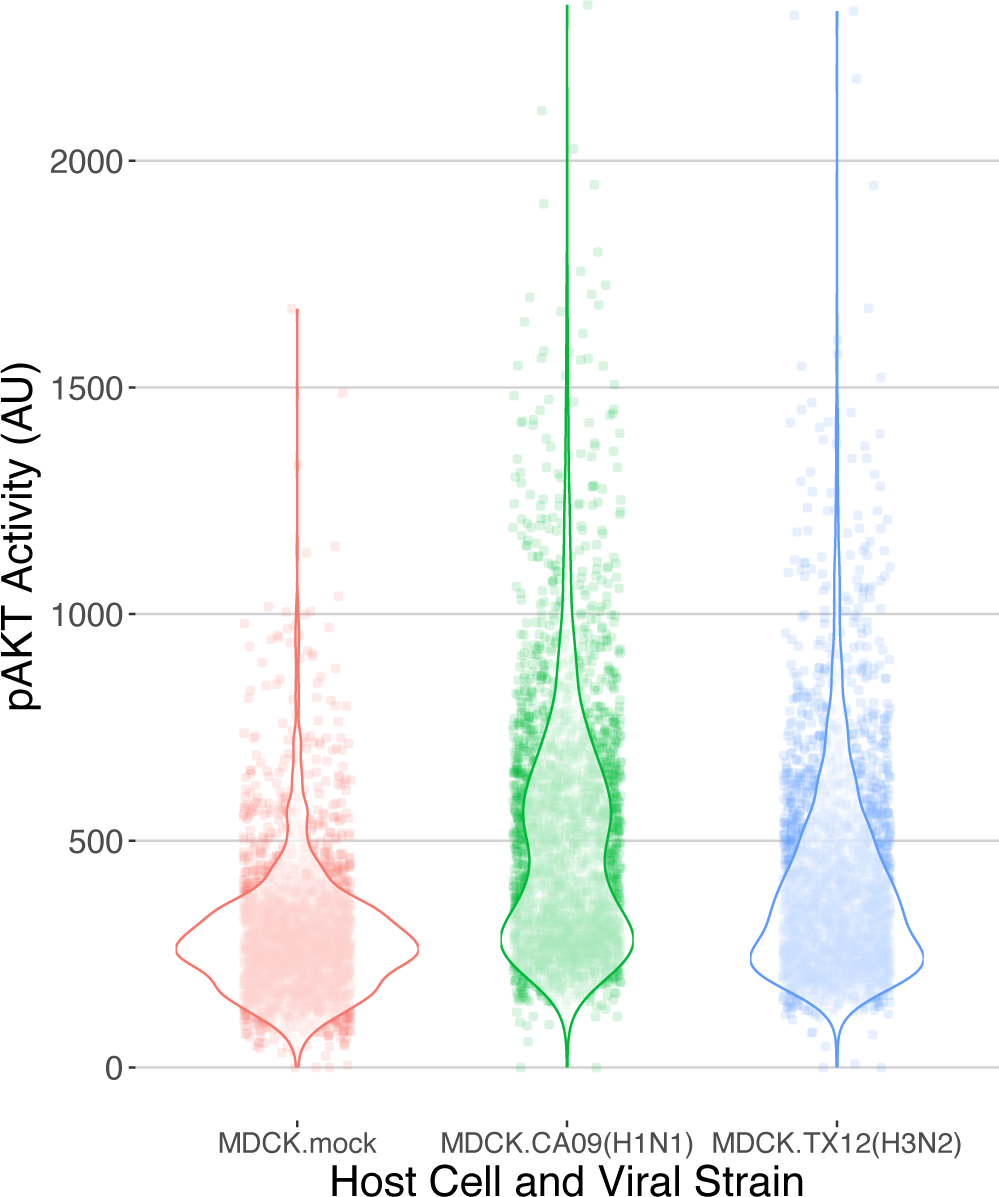
Differential activation of pAKT activity by Influenza A infection. (**A**) pAKT activity (AU) of influenza-infected MDCK-London cells. Individual points represent individual cells; a third of data points were subsampled randomly from the data set. Violin plots overlaid depict the frequency of points using density curves.

### 3. Alpelisib pre-treatment is sufficient to subvert PI3K network signal restoration by Influenza A

To determine PI3K-AKT signaling outcomes under the competing influences of influenza and alpelisib, we measured pAKT activity in cells pre-treated with increasing concentrations of alpelisib and then infected with either CA09 or TX12. We found that alpelisib completely subverted the observed viral PI3K upregulation in a dose-dependent fashion (**Figure 4, Supplementary Figure S1**); in both CA09 (Adjusted R^2^ = 0.3313, p < 0.00001) and TX12 (Adjusted R^2^ = 0.2725, p < 0.00001) strains at alpelisib concentrations from 1.25 – 40 uM. Each 1 µM increase in alpelisib led to a decrease of 6.4280 and 5.9940 AU in CA and TX infections respectively. We note that the qualitative pattern and magnitude of the decrease in AU per µM of alpelisib for cells infected with either strain was strikingly similar in uninfected cells (−4.5342 AU).

**Figure 4.**
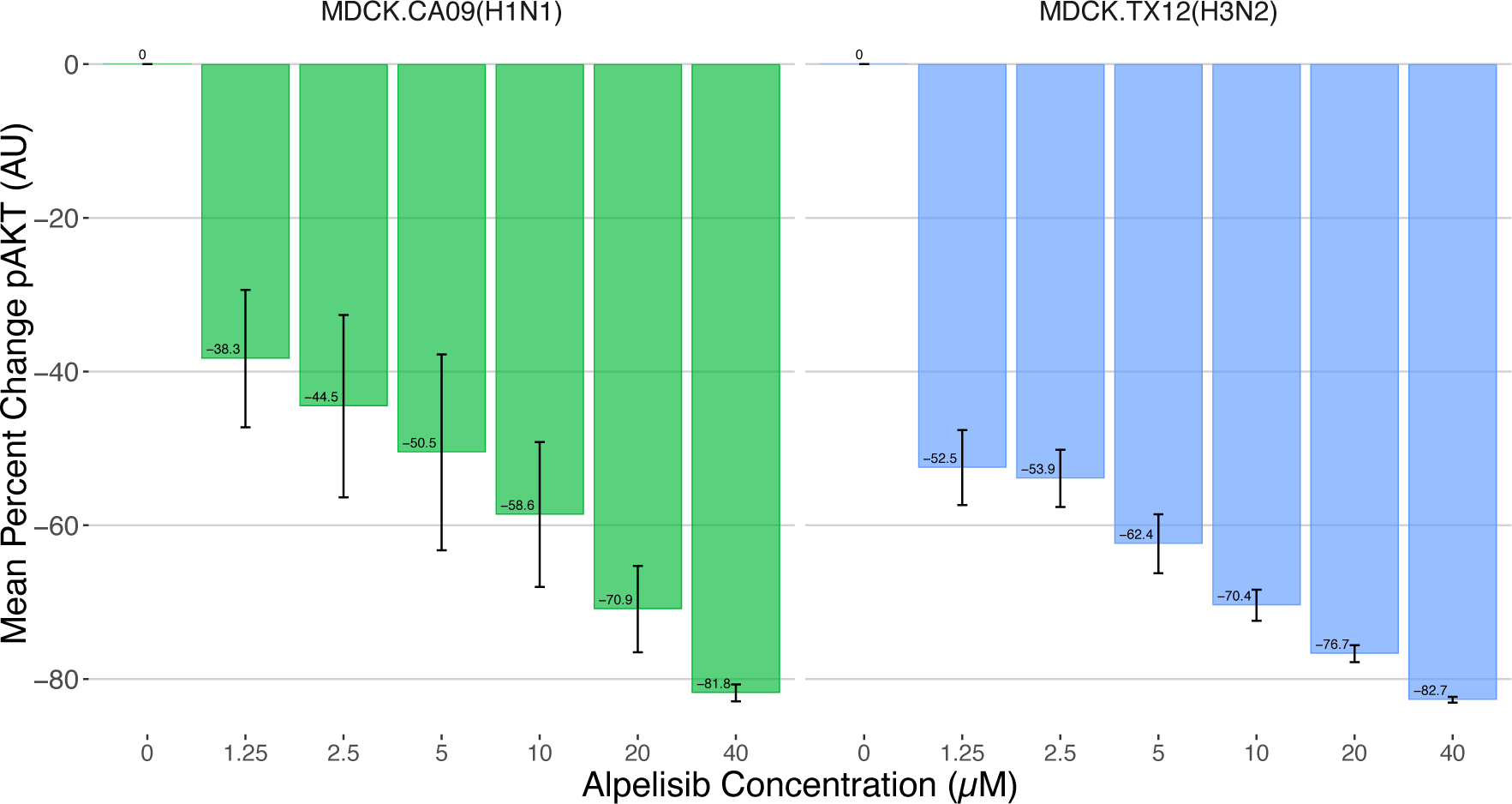
Alpelisib significantly inhibits pAKT activity during Influenza A viral infection. Mean percent change in pAKT activity (AU) of MDCK-London cells pre-treated with increasing concentrations of alpelisib and then infected with either CA09 or TX12; 0 µM alpelisib treatment group received vehicle solvent (DMSO). n = 3 bioreplicates, sem.

### 4. The Cluster-Forming Assay can titrate non-infectious/defective Influenza A particles

We developed the cluster-forming assay to simultaneously titrate fully infectious and propagation-incapable particles by combining elements of the conventional plaque assay (Cooper, 1961) and immunofocus assay (Baker, 2013). While the plaque and immunofocus assays respectively use solid or liquid overlay media to sustain inoculated monolayers for the duration of the assay, the cluster-forming assay employs a medium-viscosity overlay that remains semi-solid (Matrosovich, 2006). This medium restricts viral diffusion to neighboring cells much like a plaque assay, but can be removed for fixation, staining, and imaging of monolayers akin to the immunofocus assay (see Methods). The cluster-forming assay yields immunofluorescence (IF) images where each infection event appears as either a cluster of infected cells (productive clustering unit, PCU) or solitary infected cell foci (non-clustering units, NCU) (**Figure 5, Supplementary Figures S2-S6**). PCUs represent a productive infection mounted by a single fully infectious virus particle, whereas NCUs represent self-limiting infections mounted by propagation-incapable viral particles.

**Figure 5.**
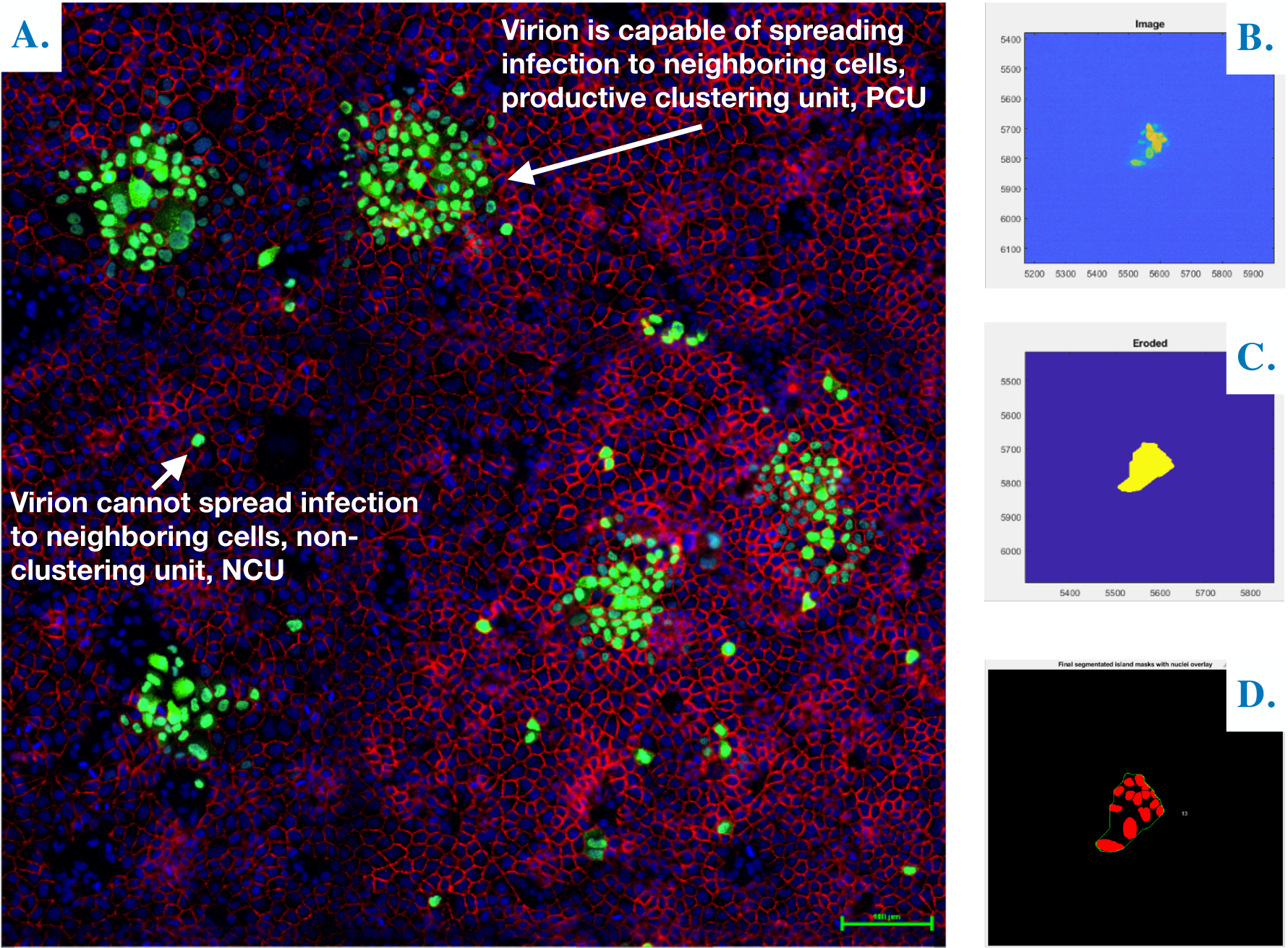
The cluster-forming assay can simultaneously titrate virions that mount productive infections and propagation-incapable virions that do not complete the infectious cycle. Influenza A Virus infection of MDCK-London cells showing productive and abortive infections. Green/GFP – A/California/07/2009 nucleoprotein; Blue/Hoechst – MDCK-London nucleus; Red/Cy5 – MDCK-London E-cadherin. **B-C**. Stepwise assembly of a mask around the nucleoprotein GFP signal in a productive clustering unit (PCU); starting from the initial cluster-forming assay IF image (**B**) down to final erosion (**C**). **D**. segmentation-mask overlay to size (i.e. number of cells) a PCU.

To count PCUs and NCUs, cluster-forming assay IF images (**Figure 5A**) were put through an automated image analysis pipeline we developed using MATLAB’s image processing toolbox. Our guiding design principle was to cordon—or mask—nucleoprotein fluorescence signals (GFP) in the IF image as independent infection events, then overlay said mask with the host nuclei segmentation Hoechst signal to reveal the number of cells each infection event had spread to. We began stepwise assembly of masks around the GFP signals (**Figure 5B-C**) by binarizing IF images with the *imbinarize* function to make object detection possible, followed by the removal of small, noisy pixels with *bwareaopen*. Masks were sequentially dilated then filled with *imdilate* and *imfill* functions respectively to smoothen them out and ensure they did not contain holes. To finish the mask assembly, masks were eroded with *imerode* to undo the signal expansion done in the dilation step (**Figure 5C**). Undesired masks were filtered out by thresholding the min/max mask area and removing masks that did not contain any nuclei, leaving bona fide infection events—or clusters—that are counted and assigned a unique identity number, a *clusterID* (**Figure 5D**).

The cluster-forming assay is a highly reproducible (**Figure 6**) improvement of the conventional immunofocus and plaque assays that provides increased resolution of viral infectivity. In addition to quantifying FIPs—as was possible with a conventional plaque assay—it is now possible to simultaneously quantify SIPs in the same sample with high-throughput. This is significant because access to two sub-populations of infectious viral particles makes it possible to determine total infectious particles, and thus relative abundances as well as; both of which are indispensable metrics for the quantitation of viral interference within the host.

**Figure 6.**
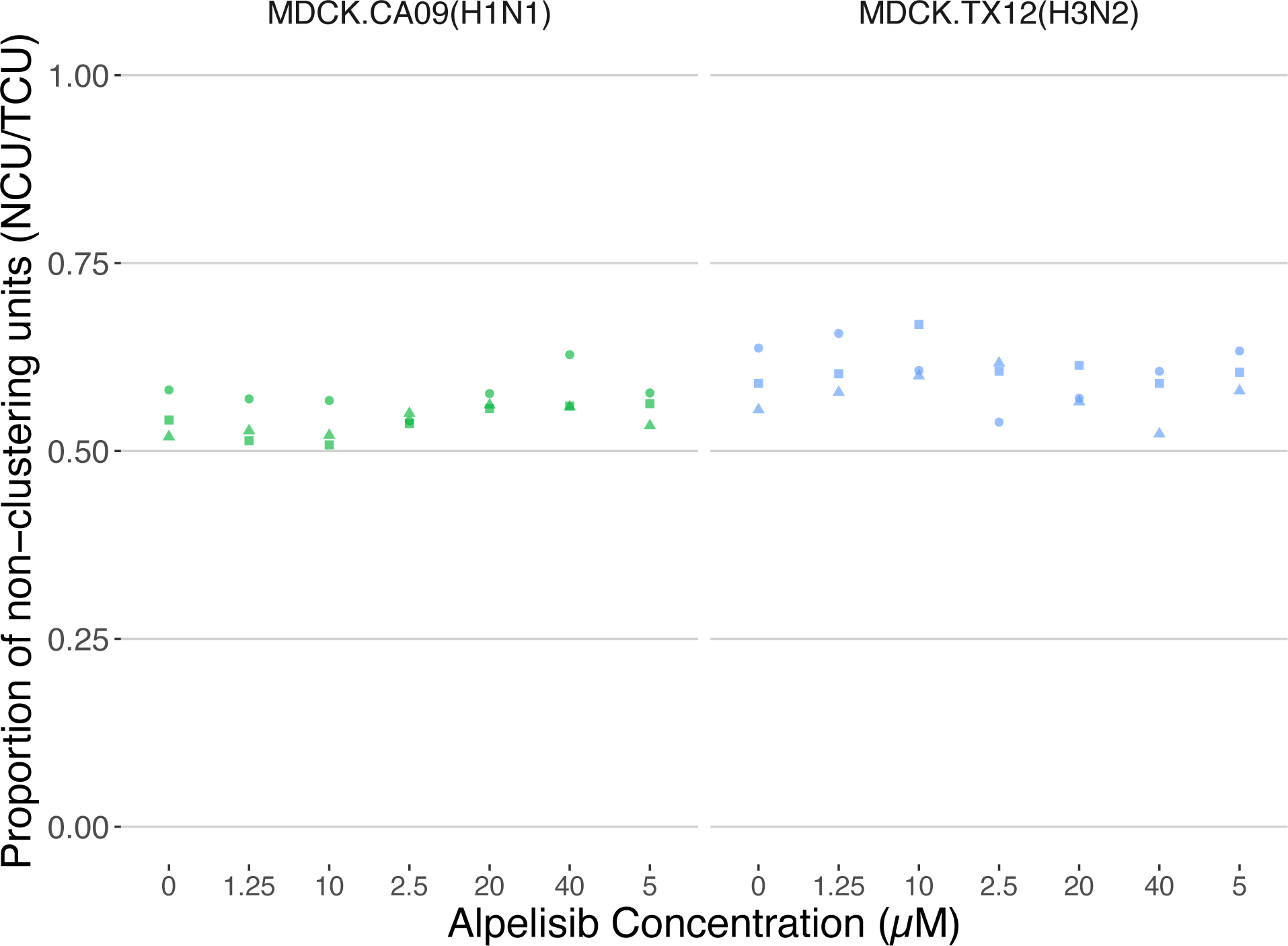
The cluster forming assay is highly reproducible. The proportion of non-clustering units (over total clustering units, aka total infection events) titrated from supernatants of 18 hr CA09 and TX12 infections of cells pre-treated with different concentrations of alpelisib. Each point is a technical replicate; i.e. a titration of the same infection supernatant. The 0 µM alpelisib treatment group received vehicle solvent (DMSO).

### 5. Alpelisib affects the production of defective particles early in Influenza A infection

To determine whether alpelisib pre-exposure of cells affected CA09 or TX12 infection, we used the cluster-forming assay to screen a broad dosage range of alpelisib pre-treatment concentrations (**Figure 2.7**).

**Figure 7.**
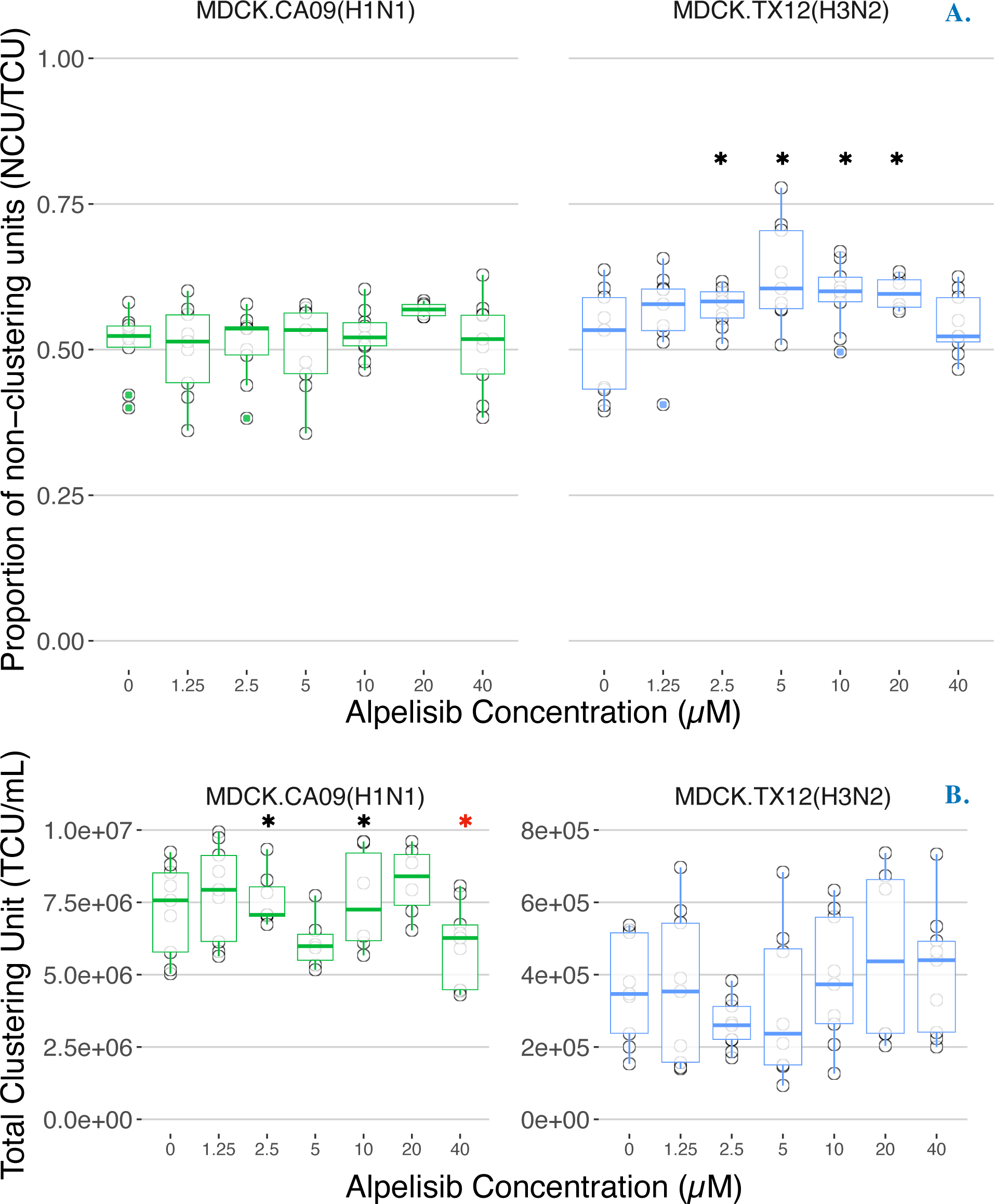
Alpelisib can affect the proportion of defective particles, as well as the total particle yield in a strain dependent manner. Proportion of (**A**) non-clustering units (NCU) and (**B**) concentration of total clustering units (TCU/mL) in CA09 and TX12 at 18 h.p.i. under different concentrations of alpelisib; no trypsin. 0 µM alpelisib treatment group received vehicle solvent (DMSO). n = 3 bioeplicates. Histogram colors denote the strain. Black asterisks denote statistically significant increases, red asterisks denote statistically significant decreases. Alpelisib had the most marked effects on the proportion of defective particles in TX12 (H3N2) (**A**, Right panel) and on total particles in CA09 (H1N1) (**B**, Left panel).

Specifically, we sought to determine if alpelisib pre-treatment (0, 1.25, 2.5, 5, 10, 20, and 40 µM) affected the production of defective/non-infectious particles, reported here as the relative abundance or proportion of NCUs. We conducted Linear mixed-effects modeling (nmle) to analyze each alpelisib treatment as a factor because a dose-dependent response was not found for either infection. Alpelisib pre-treatment significantly altered the proportion of NCUs in TX12 (p < 0.0001), but in CA09, no treatment was significantly different against the control. In TX12, concentrations of 2.5, 5, 10, and 20 µM alpelisib increased the percentage of NCUs by 6.37%, 11.946%, 8.511% and 6.70% respectively, and these effects were statistically significant (all p < 0.0388).

We also examined whether alpelisib pre-treatment affected the total viral yield, here measured by total clustering units (TCU), which are analogous to plaque forming units (PFU). In CA09 infections, alpelisib was a statistically significant factor affecting TCUs (Adjusted R^2^ = 0.8247, p < 0.00001); alpelisib concentrations of 2.5, 10, and 40 µM were significantly different from control, changing TCUs by an average of 1.21×10^6^, 1.35×10^6^, and −1.20×10^6^ TCU/mL, respectively. These values represent increases of 16.67% (2.5µM) and 18.66% (10µM), as well as a *decrease* of 16.56% in the 40µM treatment. In TX12 infections, alpelisib was not a statistically significant factor in explaining TCUs (p < 0.0632), and there were no significant differences between alpelisib treatments.

### 6. Alpelisib increases deletion-containing viral genome and total viral genome production early in Influenza A infection

We found alpelisib caused changes in infectiousness at the particle level, however the cluster forming assay does not identify the underlying genetic causes of changes in infectiousness. To specifically investigate whether alpelisib caused an increase specifically in defective viral genomes, we measured DelVGs by sequencing viral supernatants from the same infections that were used in the cluster-forming assay (see Heading 5). Two MinION flow cells yielded a total of 27.42 M reads. After quality control for well-formed amplicons (perfect 12nt UMIs, influenza A-specific terminal uni12/13 regions) and de-deduplicating unique molecular identifiers (UMI) we obtained 120,652.10 ± 70,902.90 reads per infection (CA09: 148,943 ± 78,828; TX12: 92,362 ± 49,104). We note that these read counts are produced from de-duplicated reads, reflecting an estimate of the RNA genome content whether deletion-containing or not (total viral genomes, TVG) in the supernatants.

To determine if alpelisib pre-treatment of cells increased defective viral genome production, we tested for differences in the proportion of defective viral genomes between the mock-infected control and a wide range of alpelisib concentrations. The vehicle-treated control yielded 3903.67 ± 995.55 DelVGs for CA09 and 3601.67 ± 577.42 for TX12 as detected by ViReMa, which respectively represents 2.52% and 3.07% of total viral genomes (154584, 118207). The overall proportion of defective viral genomes increased as a function of increasing alpelisib concentration in CA09 infections (Adjusted R^2^ = 0.400, p = 0.008; **Figure 2.9**, Left). Alpelisib explained 19.96% of the variation in the proportion of defective viral genomes, with each 1 µM increase of alpelisib increasing the total proportion of DVGs by 0.005929%. Infections with TX12 showed this increasing trend, however the model was not statistically significant (Adjusted R^2^ = 0.1037, p = 0.1907, **Figure 2.9**, Right).

Influenza viral genome segments have known variation in their propensity to generate defective viral genomes. In particular, the polymerase complex genes (PB2, PB1, and PA) are known to generate most of the DelVGs in a given influenza infection (Saira, 2013; Wu, 2022). Thus, made linear mixed models for each segment, separating into strain specific models, if strain was found to be a significant predictor. We examined differences in the proportion of DelVGs treating each concentration as a factor, as there was not a dose-dependent effect per-segment. We report all statistically significant results in **Table S1** (blue shading) and the full results are in the code on the GitHub Repository (https://github.com/pomoxis/alpelisib-SIP).

In CA09 infections, all statistically significant increases in DelVG proportion were found at the 20 µM pre-treatment concentration for segments PB1, PA and HA, with respective increases of 0.2473, 0.3737, and 0.0579 proportion units relative to the control mock infections. The CA09 strain has been documented to have higher DelVG production in the HA segment compared to other strains (Alnaji, 2019). For context, in mock infections polymerase segment DelVGs produced ranged from an average of 3.11% - 4.40% of total viral genomes, which increased to 24.67% - 40.48% in infections with 20µM of alpelisib. In TX12 infections, the only statistically significant changes from the control were in segments HA and M, consisting of decreases in DelVG’s of less than 1%. These changes occurred across a broader range of concentrations for the HA segment (1.25, 2.5, 5, 10 µM), than for the M segment (1.25µM).

We also analyzed the number of total viral genomes produced at different alpelisib concentrations using models for each genome segment. We found statistically significant decreases in CA09 infections at the 20µM concentration (**Figure 9A**) in all three polymerase segments and both antigenic segments. Specifically, segments PB2, PB1, and PA had decreases of −5173.3, −2371, and −6456 total viral genomes, respectively, whereas HA and NA showed decreases of −6840 and −6758 total viral genomes, respectively (**Figure 9A**; **Table 1**, gray shading). In TX12 infections, there were no statistically significant changes in TVGs according to alpelisib dose.

Collectively, these results suggest that alpelisib treatment of CA09 infections at 20 µM *increases* DelVG production and *decreases* total viral genomes in two of the polymerase complex segments (PB2, PB1) and the HA segment (**Figure 9C**), a signature of defective interference. Statistically significant changes in TX12 DelVGs were of small magnitude and total viral genome production did not show statistically significant differences across alpelisib doses. Furthermore, the overall proportion of DelVGs (regardless of segment) shows a dose-dependent increase as alpelisib concentration increased in CA09 infections (**Figure 8**). This same trend was evident in TX12 infections, but the linear regression was not statistically significant.

**Figure 8.**
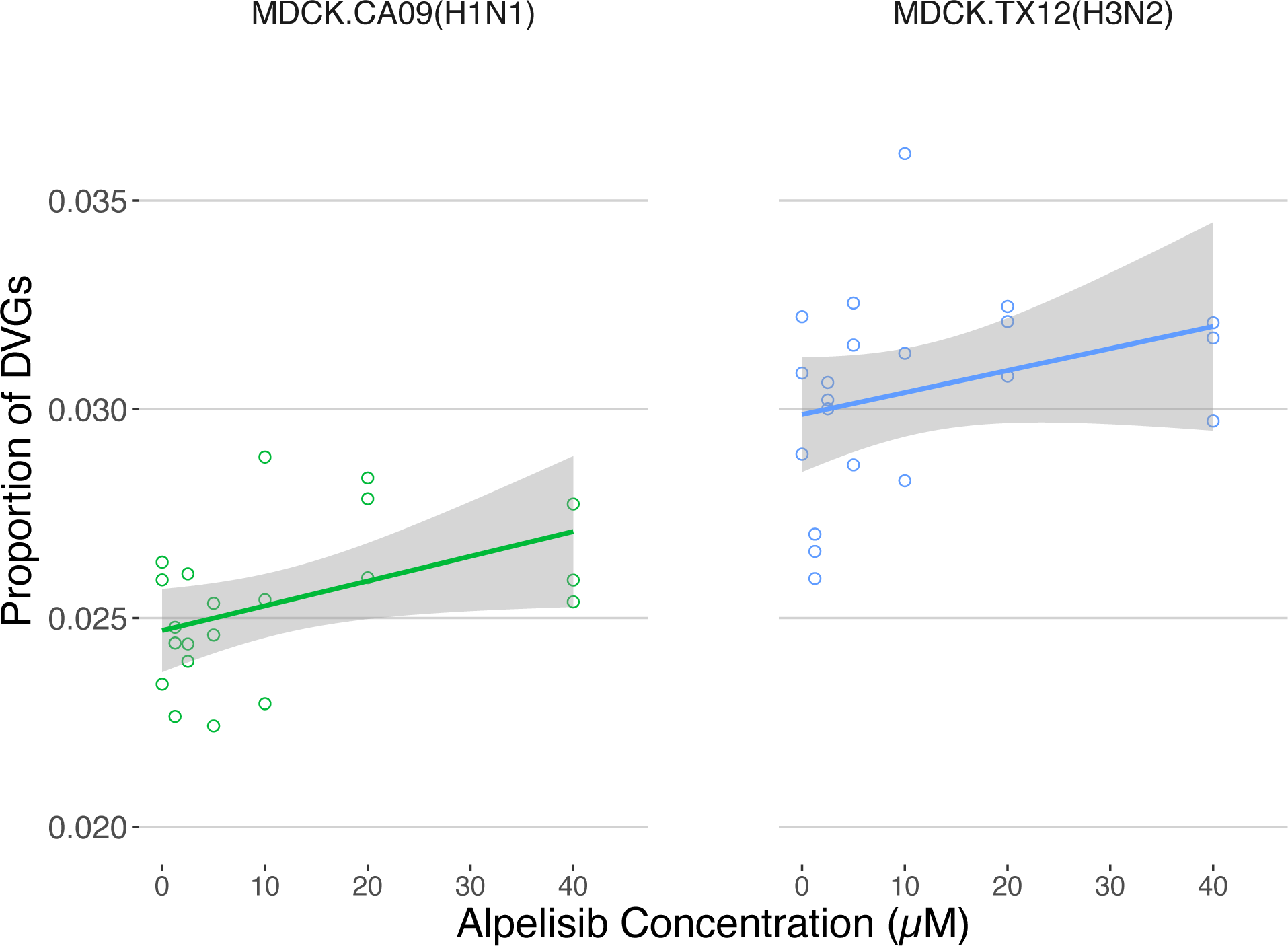
Alpelisib pre-treatment of cells increases the proportion of defective particles. Overall proportion of DelVGs (i.e. total DelVGs regardless of segment origin) as a function of concentration of alpelisib pre-treatment. The CA09 regression (Left panel) is statistically significant (p = 0.008), while TX12’s (Right panel) is not (p = 0.1907). Three independent infections with CA09 and TX12 per concentration at 18 h.p.i. under different concentrations of alpelisib pre-treatment; no trypsin. 0 µM alpelisib treatment group received vehicle solvent (DMSO).

**Figure 9.**
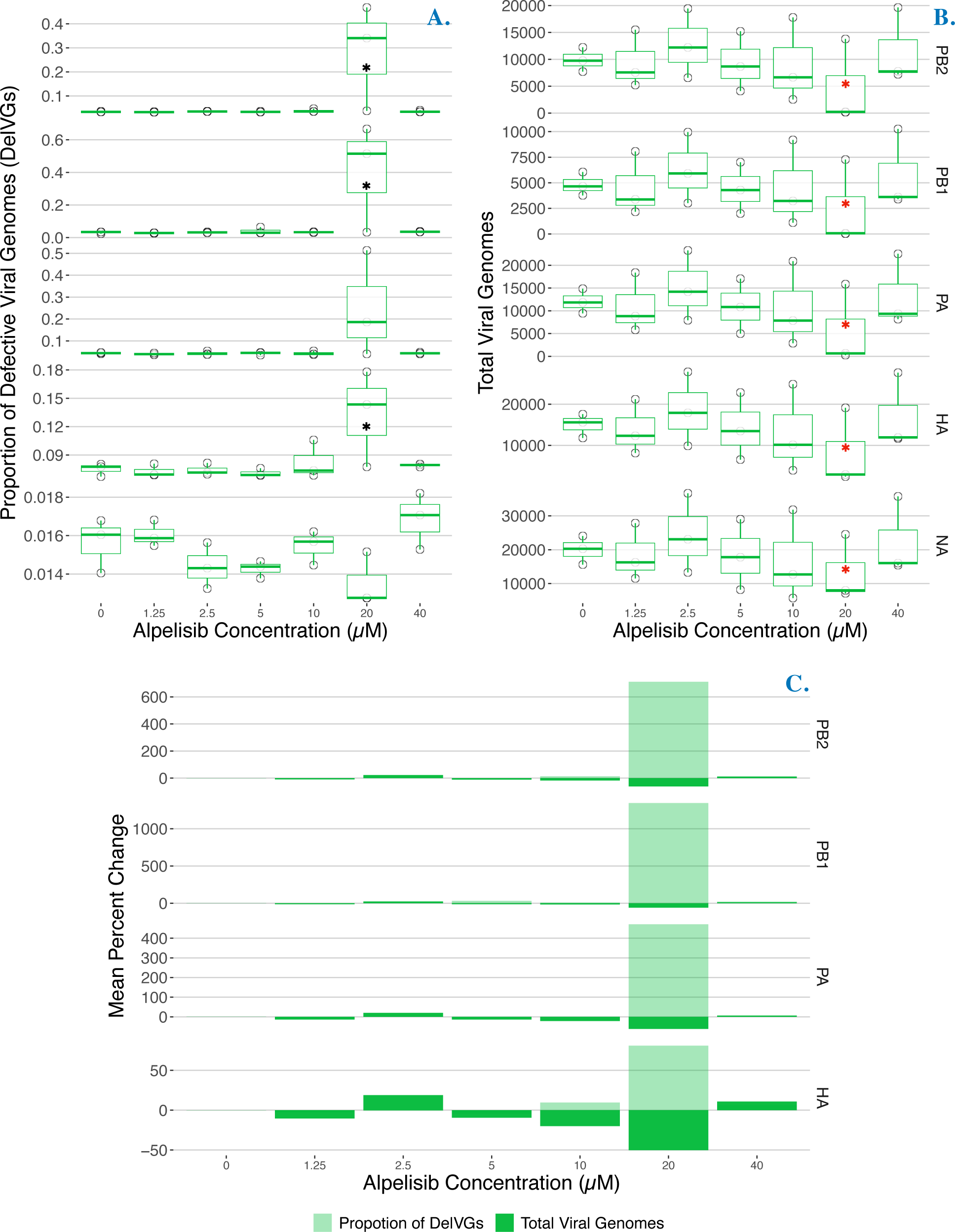
Alpelisib treatment of cells increases the proportion of deletion-containing viral genomes and decreases total viral genomes in CA09(H1N1pdm) infections. **A.** Per-segment proportion of **Del**etion-containing **V**iral **G**enomes (DelVGs) at 18 h.p.i. under different concentrations of alpelisib; no trypsin. Black asterisks indicate statistically significant increases. **B.** Per-segment **T**otal **V**iral **G**enomes (TVG) at 18 h.p.i. under different concentrations of alpelisib; no trypsin. Red asterisks indicate statistically significant decreases. **C.** Average percentage change in **Del**etion-containing **V**iral **G**enomes (DelVGs, light green) and **T**otal **V**iral **G**enomes (TVG, dark green) under different concentrations of alpelisib. The 0 µM alpelisib treatment group received vehicle solvent (DMSO). n = 3 bioreplicates, sem.

### 7. Alpelisib increases average deletion size in the polymerase segments

We found Alpelisib increased the number of DelVGs, and wanted to examine how many different DelVGs were formed and distribution of DelVGs also changed. To meet this goal, we used ViReMa to enumerate all the unique species of DelVGs in each segment and used linear mixed models to test for differences among alpelisib concentrations; that is we examined how many different DelVGs were formed, but did not evaluate their abundance in these analyses. Alpelisib increased the average deletion size of polymerase complex segment DelVGs species in both the H1N1 and H3N2 strain (**Figure 10**, **Table S2**). Specifically, CA09 infections with 20µM of alpelisib had increases in average deletion size on all three polymerase segments (PB2: 124.00 bps; PB1: 193.78 bps; PA: 148.511 bps) and the HA segment (52.846 bps). TX12 registered more modest increases in average deletion size in the PB2 (20µM: 61.96 bps) and PB1 (20µM: 73.45 bps; 40µM: 78.46) segments. Additionally, CA09 also registered *decreases* in average PB2 deletion sizes ranging from 64.287 - 85.754 bps at the 1.25, 2.5, and 5µM concentrations.

**Figure 10.**
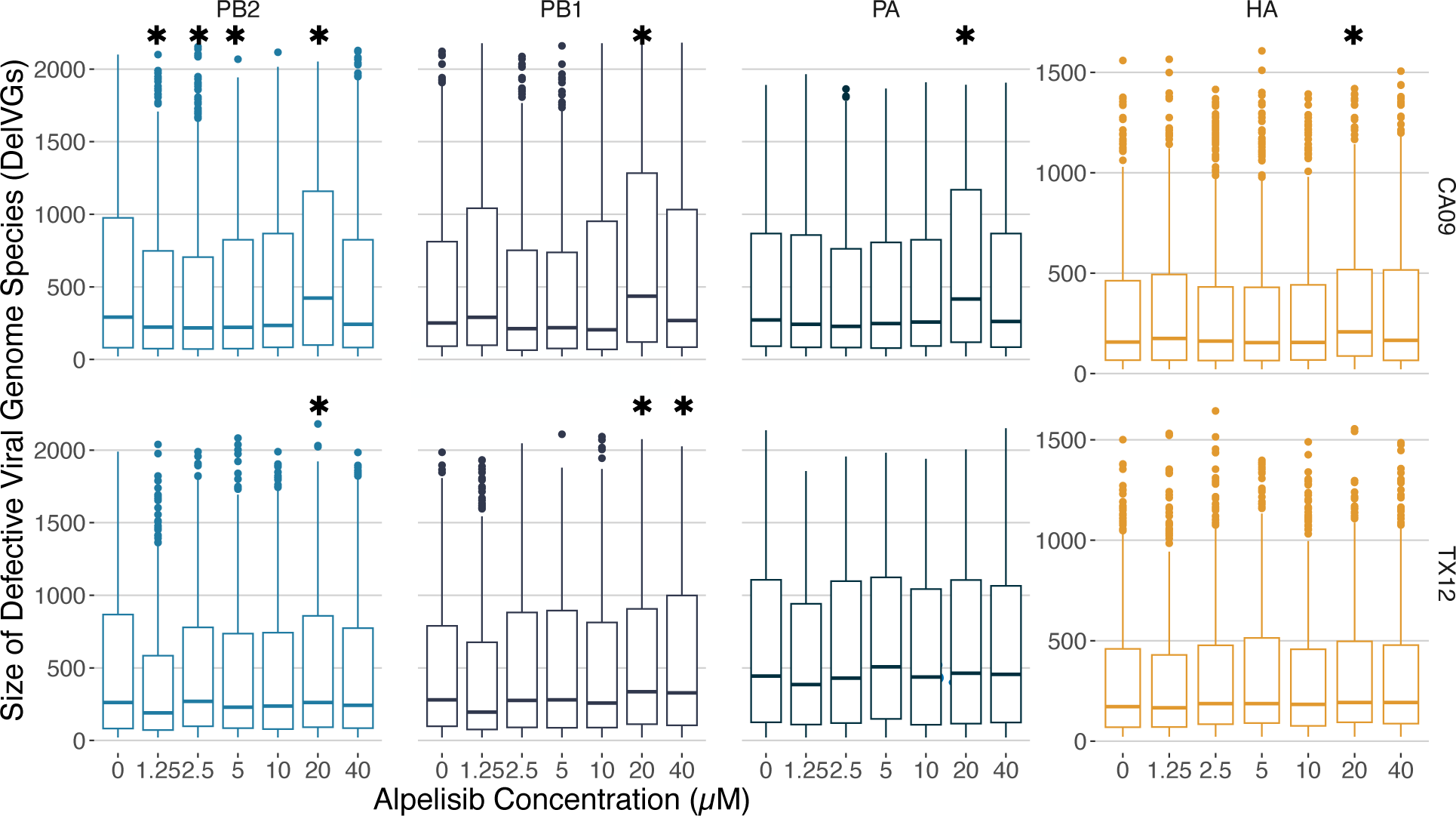
Alpelisib increases average deletion size in the polymerase segments. Size (in basepairs, bps) distribution of distinct deletions in defective viral genomes (here measured **Del**etion-containing **V**iral **G**enomes (DelVGs) produced by CA09 (H1N1) and TX12 (H3N2) infections under different concentrations of Alpelisib. Asterisks indicate statistically significant changes in mean size compared to the mock infection. Top row: CA09 (H1N1) strain; bottom row: TX12 (H3N2) strain. Box-plot colors represent different genome segments, note the different y-axis scale in segment HA plot.

The change in the average deletion size in the DelVG population appeared to be due to an overall decrease of DelVGs with small deletions and most of the remaining DelVGs having larger deletions (**Figure 10**).

## Discussion

To investigate the dependence of influenza progeny infectivity on host cell metabolic signaling, we reprogrammed host PI3K network signaling during flu infection with alpelisib, monitored the intervening metabolic state with cellular-level resolution immunofluorescence microscopy, and determined the change in proportion of non-infectious progeny virus with our newly developed cluster-forming assay. The proportion of non-infectious particles in a flu infection is emerging as a crucial determinant of pathogenic outcomes, and non-infectious particles are being directly used as antiviral treatments (Smith, 2016; Meng, 2017; Wasik, 2018; Zhao, 2018; Bdier, 2019; Yamagata, 2019; Tapia, 2019; Harding, 2019). We established that host cell PI3K network signaling activity can influence the proportion of non-infective particles produced by influenza in two strains. First, we found that alpelisib suppresses PI3K-AKT pathway signaling activity in MDCK-London cells in a dose-dependent manner (**Figure 1A**). Second, we found that alpelisib treatment keeps PI3K-AKT signaling pathway activity suppressed during Influenza A infection in MDCK-London cells (**Figure 4**), counteracting influenza’s upregulation of PI3K signaling (**Figure 3**; Hale, 2006; Ayllon, 2012). As predicted by our hypothesis, we found that alpelisib treatment induces a reproducible and statistically significant increase in the proportion of non-infectious progeny during Influenza A infection in the TX12 strain (**Figure 7**). We also found that that there is a statistically significant dose-dependent increase in the overall proportion of defective viral genomes (**Figure 8**) in CA09 infections. Moreover, we found that alpelisib increases the proportion of deletion-containing viral genomes produced by polymerase complex and antigenic segments in CA09 infections (**Figure 9A**) and that these increases are coincident with decreases in total viral genomes (**Figure 9B-C**), suggesting DelVGs were acting as defective interfering genomes. Finally, we documented that alpelisib affected average deletion size of polymerase complex segment DelVGs in both CA09 and TX12 strains (**Figure 10**). Collectively, these findings establish the host cell’s metabolic signaling profile as a means to directly modulate the infectivity of progeny Influenza A virions and DelVGs, further validating the use of metabolic signal modulation as a means to drive influenza infections toward milder clinical outcomes.

To our knowledge, our study is the first to examine the role of host metabolic state in the production of non-infectious influenza progeny particles and defective viral genomes (Dimmock, 2014; Manzoni, 2018; Vignuzi, 2019; Wu, 2022). Specifically, we show differential dysregulation of PI3K signaling activity in a non-tumorigenic cell line during infection by two different Influenza A strains, CA09 and TX12, then override this host-virus interaction using alpelisib, and finally show an increase in DelVGs and defective particle production. The host metabolic state can affect influenza infection in terms of clinical outcomes as shown in studies of obesity and cancer. For instance, obesity—a host metabolic state characterized by chronic inflammation and dysregulated immune responses—has been associated with increased titers of infectious progeny (Honce, 2019; Honce, 2020). Similarly, at the cellular level, cross-talk between Influenza A and host metabolic signaling effectors has been shown to affect the production of infectious progeny (Hale, 2006; Li, 2008; Smallwood, 2017; Kuss-Duerkop 2017). However, the production of non-infectious progeny during different metabolic states had not previously been investigated; an extremely relevant line of investigation given that up to 90% of total viral particles are non-infectious (Brooke, 2013; Brooke, 2017; Diefenbacher, 2018), and that these particles have a role in clinical outcomes (Dimmock, 2014; Vasilijevic, 2017). Our findings confirm that interrupting virus-induced upregulation of host growth signaling can increase non-infectious Influenza A particle production, providing novel insight into the crosstalk between Influenza A and host metabolism. The molecular mechanisms that underlie and influence non-infectious particle formation remain unclear. Until now, these investigations have focused primarily on the viral side of the equation, identifying viral genome mutations and infection multiplicity as variables influencing the production of non-infectious progeny across different strains (Von Magnus, 1954; Rodriguez, 2013; Vasilijevic, 2017; Perez-Cidoncha, 2014). Our study addresses the need to better understand the still unknown mechanisms that spawn non-infectious progeny by providing a more complete picture of how metabolic state affects Influenza A pathogenesis.

Our study combines single-cell immunofluorescence quantification with an automated assay that quantifies the proportion of non-infectious virus particles, providing a more accurate measurement of the infectious potential of a virus population (Brooke, 2013). Building on previous methods (Brooke, 2013; Amarilla, 2021; Cacciabue, 2019), our cluster-forming assay combined the infection localization of a conventional plaque assay with the immunocytochemical staining and microscopy of the standard immunofocus assay. By pairing this assay with an automated image analysis pipeline, we were able to capture influenza infectivity at a more detailed resolution than is possible with either parent assay alone (See Supplementary Material). By resolving NCUs (non-infectious particles) and PCUs (fully infectious particles) apart from each other, the cluster-forming assay has revealed strain-specific and dose-specific effects of alpelisib on key markers of defective interference at just 18 h.p.i. These novel outcomes represent early onset alterations to the trajectories of standard CA09 and TX12 infections, and each of these altered trajectories is uniquely desirable for different real-world therapeutic and prophylactic applications, provided they persist into later time points. Pharmacologically increasing the *in situ* spawn rate of non-infectious particles is a viable and novel therapeutic possibility, provided the underlying physiological factors become better understood. One aspect that our study does not address is the definition of the non-infectious particle component. Although the cluster-forming assay accurately titrates infectious and non-infectious particles, it was not designed to identify whether those non-infectious particles represent DIPs, particles with lethal or nonsense mutations, or particles that contain segments with defects in transcription (Brooke 2013); an area that requires further study. We additionally expect the cluster-forming assay to facilitate future screens to uncover evermore druggable modulators of *in situ* non-infectious particles and DelVG production during Influenza A infection.

Based on the established cross-talk between effector proteins of both host cell metabolic signaling and Influenza A (Hale, 2006; Li, 2008; Smallwood, 2017; Kuss-Duerkop 2017), we hypothesized that progrowth metabolic signal inhibition with alpelisib would induce abortive infectivity in progeny flu particles and increase DelVGs. Our predictions proved out, and uncovering more inducers in this manner will guide future investigations into the molecular mechanism through which DelVGs emerge and non-infectious progeny particles accumulate. These findings suggest that the host context can shift the makeup of the DelVG population, shaping the potential virus-virus interactions (Leeks et al., 2023; Díaz-Muñoz et al., 2017) altering the course of the infection. Knowledge of these mechanisms will facilitate the development of more targeted abortive infectivity induction strategies for broad-spectrum anti-influenza therapeutics. DI is already being weaponized in the form of exogenously administered recombinant Influenza A virions called therapeutic interfering particles (TIPs), which have been engineered to contain one or more DelVGs. TIPs are propagation-incapable, and their administration attenuates Influenza A pathogenesis in a strain-indiscriminate (Smith 2016; Zhao, 2018), dose-dependent manner (Smith, 2016; Meng, 2017; Wasik, 2018; Zhao, 2018; Bdier, 2019; Yamagata, 2019; Tapia, 2019; Harding, 2019). Our study opens the possibility of using host cell metabolic state as a strategic therapeutic target because of its readily responsive and reversible system-wide reach. Picomolar perturbations of host cell metabolism can drive system-wide reconfiguration of critical processes into countless unique endpoints; too many endpoints for Influenza A to possibly adapt against. Future research should address the composition of non-infectious progeny particles, and which PI3K downstream effector pathways transduce the signal(s) that ultimately impacts *de novo* non-infectious particle emergence. In sum, our research shows the promise of the host cell’s vast metabolic signaling network as a quick-response, therapeutically actionable, druggable target with the potential to steer flu pathology away from fatal towards mild outcomes.

## Acknowledgements

This project was funded by NIH grants 4R00AI119401-02 and 1R01AI179873-01 to SDM. IA was supported by NIH grant 4R00AI119401-02 and 1R01AI179873-01 to SDM and a Craft Consult Biotechnology Dissertation Fellowship. We thank members of the Albeck lab for kindly provided training and access to his fluorescence microscopy resources and infrastructure. Enoch Baldwin provided helpful feedback on this manuscript. Ted Ross kindly provided strains from his collection.

