## Supplementary Materials for "Influenza A defective viral genomes and non-infectious particles are increased by host PI3K inhibition via anti-cancer drug alpelisib"

### Extended Results

#### *Insulin activates PI3K network signaling in MDCK-London cells*

In addition to exposing uninfected and virus-infected MDCK-London cells to increasing concentrations of alpelisib, a parallel experiment was run in which the same test groups were co-exposed to 10ug/mL of insulin. This experiment confirmed the potentiating effect of insulin on PI3K-AKT signaling activation (positive control for pAKT activation). pAKT activation levels in all parallel insulin exposed experiment groups are higher than their no-insulin counterparts (**Figure S1**).

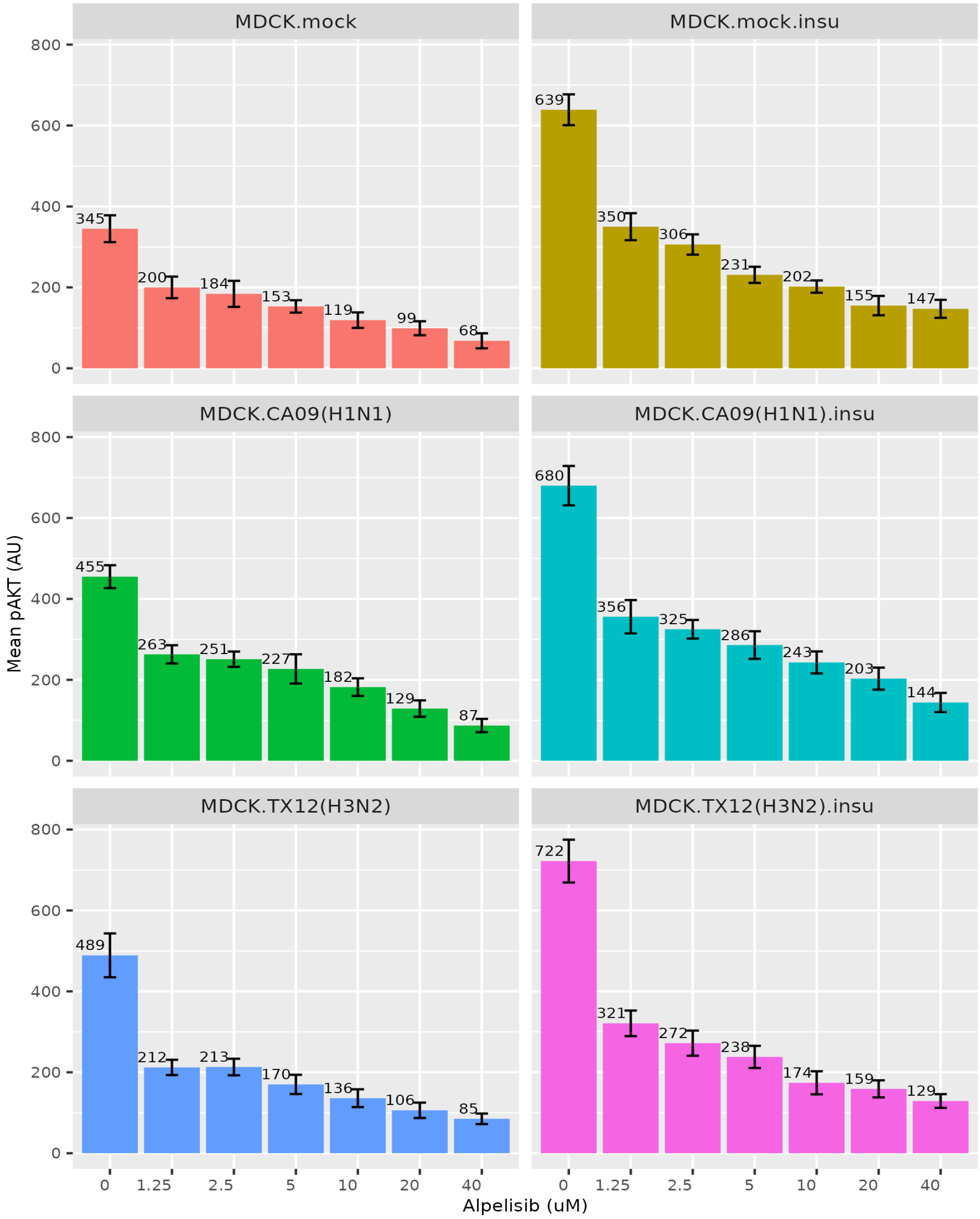


**Supplementary Figure S1.** pAKT activation in mock- and virus-infected MDCK-London cells that were either exposed to different concentrations of alpelisib (left), or co-exposed to 10ug/mL of insulin in addition to alpelisib (right). 0 µM alpelisib treatment group received vehicle solvent (DMSO). n = 3 bioreplicates, sem.

| Strain | Measure | Segment | Alpelisib Treatment (µM) | Estimate | Std. Error | t value | Pr(>\|t\|) | |
|---|---|---|---|---|---|---|---|---|
| **CA09** | PropDVG | PB2 | 20 | 0.2473223 | 0.0679473 | 3.64 | 0.00339 | ** |
| **CA09** | PropDVG | PB1 | 20 | 0.373685 | 0.102129 | 3.659 | 0.00327 | ** |
| **CA09** | PropDVG | HA | 20 | 0.0578874 | 0.0168425 | 3.437 | 0.00492 | ** |
| **TX12** | PropDVG | HA | 1.25 | -0.0060637 | 0.0008546 | -7.095 | 1.26E-05 | *** |
| **TX12** | PropDVG | HA | 2.5 | -0.0024982 | 0.0008546 | -2.923 | 1.28E-02 | * |
| **TX12** | PropDVG | HA | 5 | -0.0043145 | 0.0008546 | -5.048 | 0.000285 | *** |
| **TX12** | PropDVG | HA | 10 | -0.0026667 | 0.0008546 | -3.12 | 0.00885 | ** |
| **TX12** | PropDVG | M | 1.25 | -0.0048573 | 0.0011397 | -4.262 | 0.0011 | ** |
| **CA09** | TVG | PB2 | 20 | -5173.3 | 1893.1 | -2.733 | 1.82E-02 | * |
| **CA09** | TVG | PB1 | 20 | -2371 | 1040.4 | -2.279 | 4.18E-02 | * |
| **CA09** | TVG | PA | 20 | -6456 | 2235 | -2.889 | 1.36E-02 | * |
| **CA09** | TVG | HA | 20 | -6840 | 2576 | -2.655 | 0.02099 | * |
| **CA09** | TVG | NA | 20 | -6758 | 2921 | -2.314 | 0.039197 | * |

Supplementary Table S1. **Alpelisib affects the proportion of deletion containing viral genomes and total viral genomes produced in infections**. Statistically significant predictors of the average size of of **Del**etion-containing **V**iral **G**enomes (DelVGs, blue shading) and and Total Viral Genomes (TVG, gray shading) including their parameter estimates from per-segment linear mixed models.

| Strain | Measure | Segment | Alpelisib Treatment (µM) | Estimate | Std. Error | t value | Pr(>\|t\|) | |
|---|---|---|---|---|---|---|---|---|
| **CA09** | size (bp) | PB2 | 1.25 | -69.648 | 28.061 | -2.482 | 1.31E-02 | * |
| **CA09** | size (bp) | PB2 | 2.5 | -85.754 | 25.689 | -3.338 | 8.49E-04 | *** |
| **CA09** | size (bp) | PB2 | 5 | -64.287 | 27.952 | -2.3 | 2.15E-02 | * |
| **CA09** | size (bp) | PB2 | 20 | 124.005 | 31.561 | 3.929 | 8.63E-05 | *** |
| **TX12** | size (bp) | PB2 | 20 | 61.96 | 30.2 | 2.052 | 0.0402 | * |
| **CA09** | size (bp) | PB1 | 20 | 193.782 | 49.145 | 3.943 | 8.23E-05 | *** |
| **TX12** | size (bp) | PB1 | 20 | 73.45 | 32.35 | 2.27 | 0.0232 | * |
| **TX12** | size (bp) | PB1 | 40 | 78.46 | 32.3 | 2.429 | 0.0152 | * |
| **CA09** | size (bp) | PA | 20 | 148.511 | 23.739 | 6.256 | 4.15E-10 | *** |
| **CA09** | size (bp) | HA | 20 | 52.846 | 18.431 | 2.867 | 4.16E-03 | ** |

Supplementary Table S2. **Alpelisib affects average deletion size in infections**. Statistically significant predictors of the average size of of **Del**etion-containing **V**iral **G**enomes and their parameter estimates from per-segment linear mixed models.

### Extended Methods

#### *Cluster-forming Assay*

The cluster-forming assay combines the infection localization of a conventional plaque assay with the immunofluorescence (IF) staining and microscopy of a conventional immunofocus assay to capture influenza infectivity at a deeper resolution than possible with either parent assay alone.

Overnight MDCK-London cells—1.04 x $10^5$ total cells per well—were seeded into collagen-treated, glass-bottom 96-well tissue culture plates in MEM plus 5% FBS media to achieve 100% confluence in 24 hr. Confluent monolayers were then inoculated with serial dilutions of virus stock and incubated for 1 hr to allow for virus-monolayer adsorption, after which inoculum was aspirated and monolayers washed with MEM plus 2% bovine serum albumin and 1% Anti-Anti (Virus Infection Media; VIM). At this juncture, the conventional plaque assay or immunofocus assay would respectively see a solid or liquid overlay medium applied to the inoculated monolayers. The cluster-forming assay, on the other hand, applies a low to medium-viscosity overlay medium (VIM plus 4% carboxymethyl cellulose and 1 ug/mL TPCK-Trypsin) that remains in a semi-solid state at the end of the assay. This viscous overlay restricts diffusion of progeny virus to directly adjacent cells much like a conventional plaque assay, but has the added benefit of being removable via an aspirator pipette so that monolayers may be fixed, stained with IF antibodies, and imaged. Overlaid monolayers were incubated an additional 11 hours, at which point overlay media was aspirated and monolayers fixed with 4% PFA. Fixed monolayers were stained with fluorophore-conjugated IF antibodies targeting Influenza A nucleoprotein, counterstained with Hoechst, and imaged via fluorescence microscopy. The output at this juncture is an IF image of an flu-infected monolayer (**Supplementary Figure 2.1**) wherein each cluster of infected cells represents a productive infection mounted by a single propagation-capable virion—a *productive clustering unit* (PCU)—while solitary infection foci represent abortive infections mounted by propagation-incapable virus—*non-clustering units* (NCU).

**2.1A**

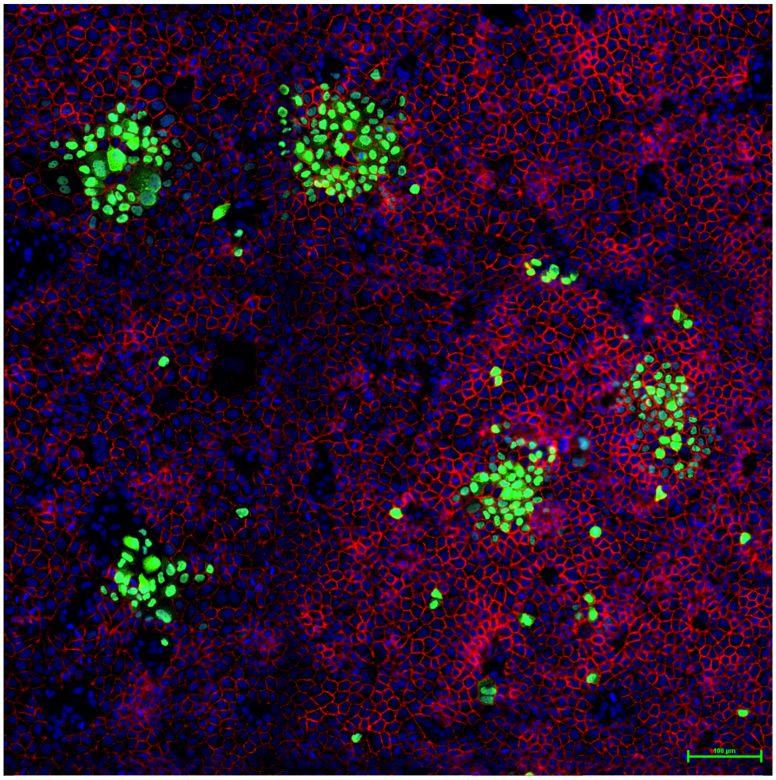

**2.1B**

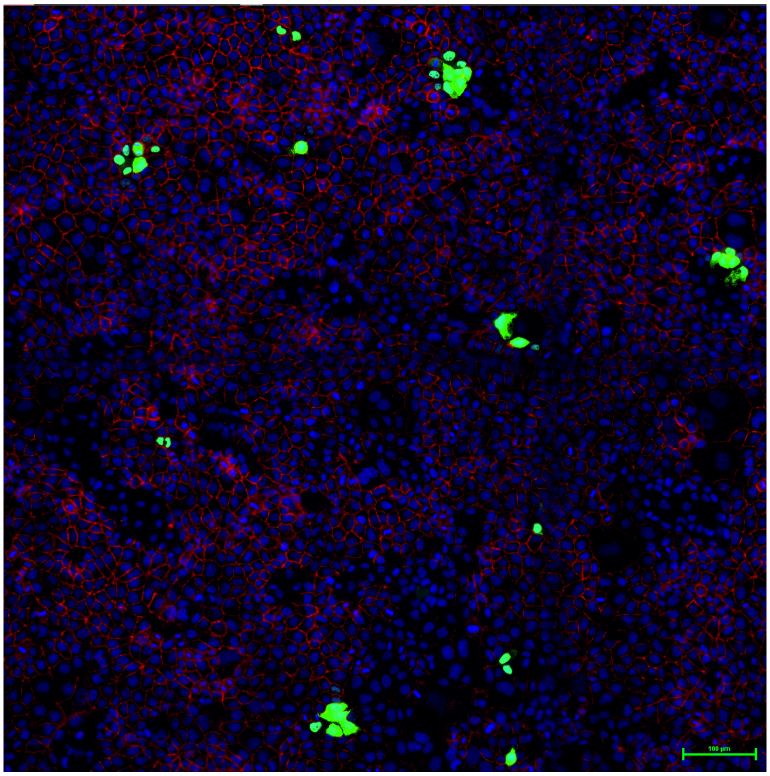


Supplementary Figure S2.1. Cluster-forming assay of Influenza A Virus on MDCK-London cells showing productive and abortive infections. Green/GFP – A/California/07/2009 nucleoprotein (**A**), A/Texas/50/2012 nucleoprotein (**B**); Blue/Hoechst – MDCK-London nucleus; Red/Cy5 – MDCK-London E-cadherin (**A**) or β-catenin (**B**).

Our proof of concept cluster-forming assays worked as designed; propagation-incapable virus mounted abortive infections as evidenced by NCUs, while propagation-capable virus mounted productive infections as evidenced by PCUs. Ordinarily, PCUs and NCUs would be tallied and titrated, but minor optimization of a few parameters was necessary to boost assay precision. Chief among these parameters were *monolayer integrity* and *PCU spillover*.

**Monolayer Integrity**: Monolayer damage and stripping undermines cluster-forming assay precision because the signal of an infection event is diminished—or lost outright—with each sloughed cell. This is especially relevant for abortive infections, whose assay signal is transmitted by a single cell, the sloughing of which results in underreporting and underestimation of NCU titer and NCU proportion. Therefore, it is imperative that monolayer integrity be preserved at all steps. We discovered that not prewarming reagents to room temperature or 37 °C—i.e. rapid reduction in temperature—caused monolayers to peel (**Supplementary Figure 2.2A**). That said, monolayer damage was consistently and primarily observed in monolayer regions of lower cell density (**Supplementary Figure 2.2B**). We found that seeding at higher cell density—upwards of 1.6 x $10^4$ to 1.04 x $10^5$ cells per well—and seeding more evenly with a wide-bore 2mL serological pipette in place of a multichannel pipette was sufficient to mitigate monolayer damage in future assays (**Supplementary Figure 2.3A**). However, PCU spillover and streaks remained an issue; especially for a fast growing strain like the pandemic A/California/07/2009(H1N1) (**Supplementary Figure 2.3B-C**).

**2.2A**

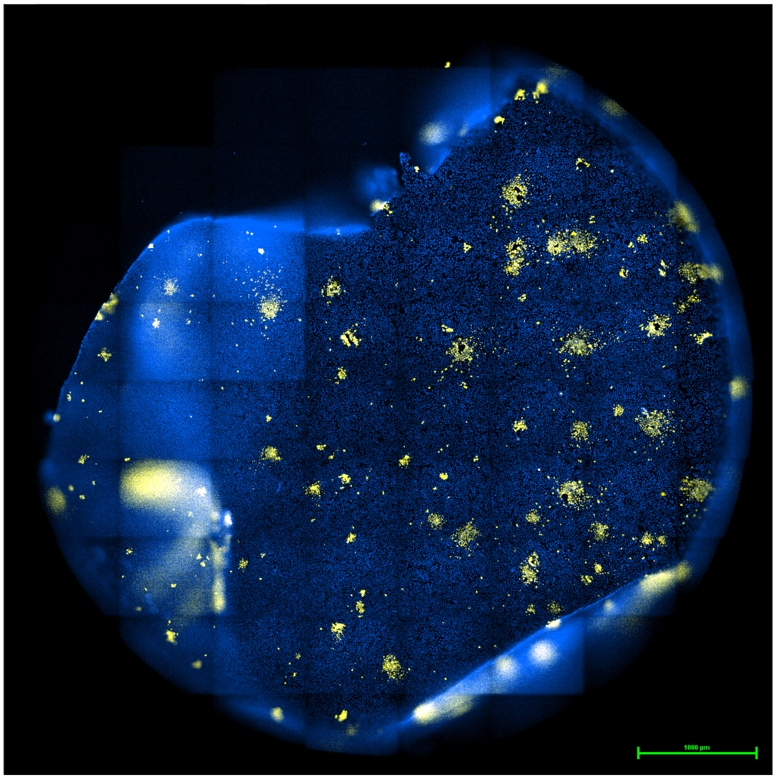


**2.2B**

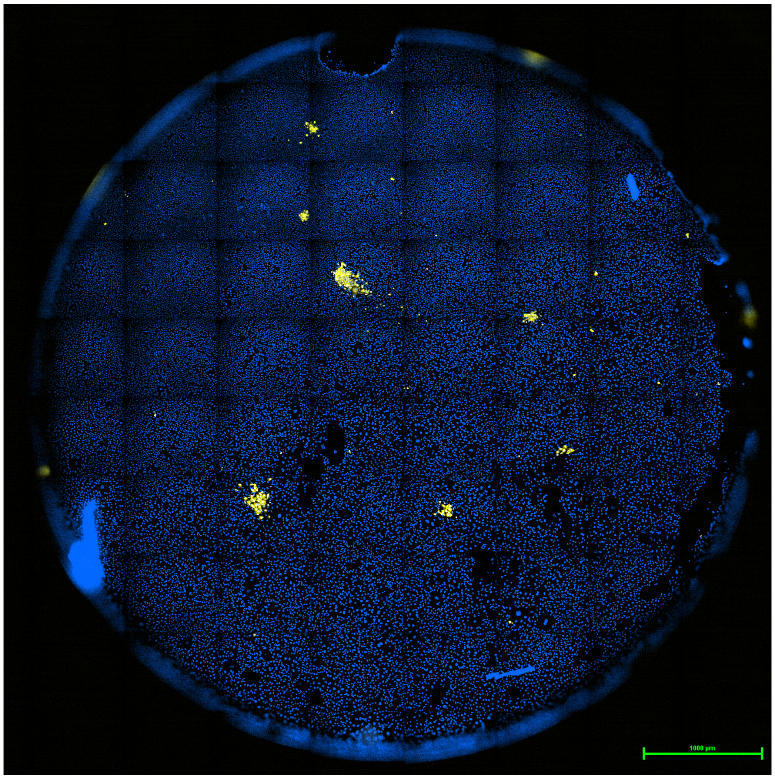

**2.2C**

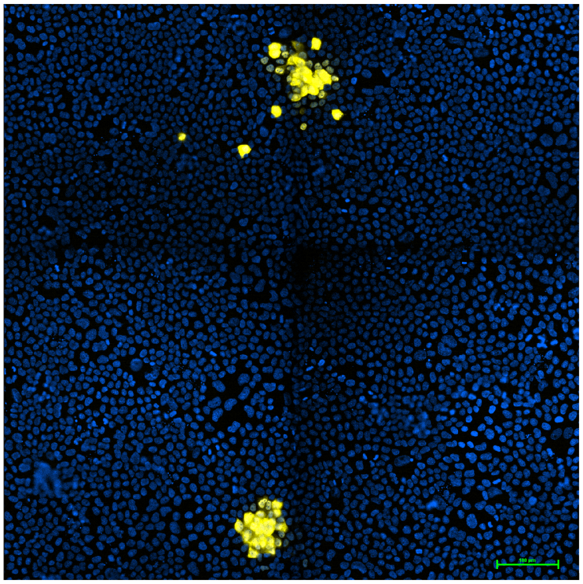

**2.2D**

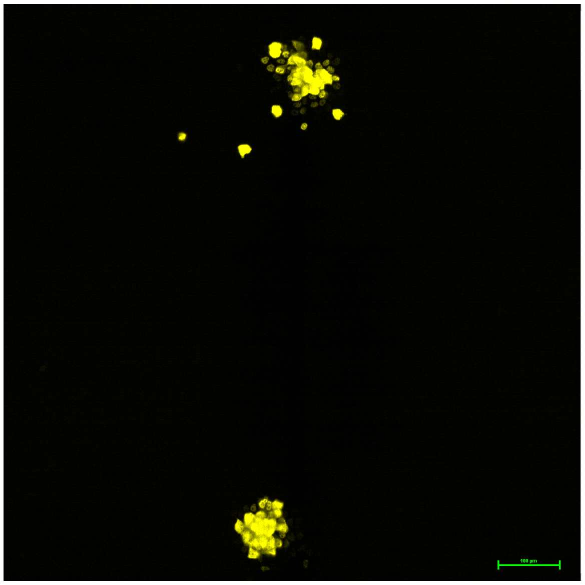

**2.2E**

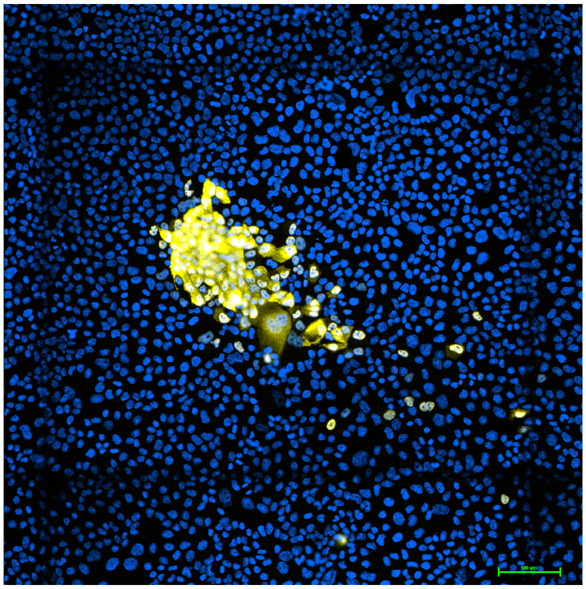

**2.2F**

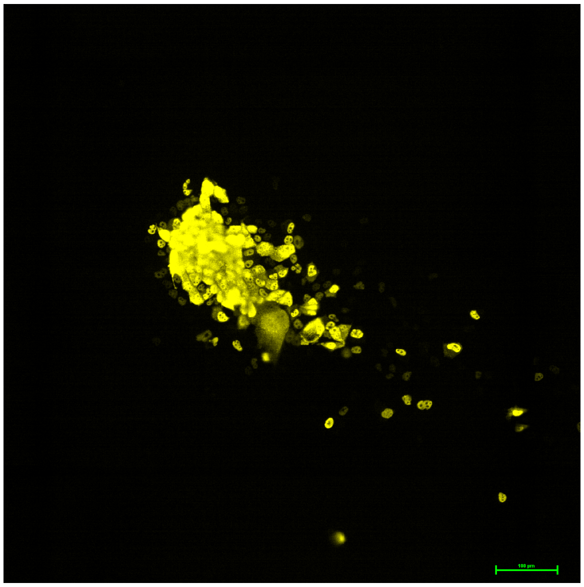

Supplementary Figure S2.2. Early proof of concept cluster-forming assay. (**A**) Monolayer peeling following exposure to cold (4 °C) reagents; always pre-warm reagents to between room temp and 37 °C before use. (**B**) Monolayer damage in low cell density areas; seed monolayers uniformly and at sufficient cell density. (**C, D**) PCU spillover. (**E, F**) PCU streak/comet. Yellow/YFP – A/Texas/50/2012 nucleoprotein ; Blue/Hoechst – MDCK-London nucleus.

**PCU spillover**: In a cluster-forming assay, *spillover* has occurred if there are satellite single-cell infection event(s) surrounding a PCU (**Supplementary Figure 2.2C-F, 2.3B-C**). Spillover undermines assay accuracy because it is unclear if these solitary infection events are progeny virus spawned by the closeby PCU, or a bona fide NCU from the initial inoculum. To mitigate PCU spillover, we pursued optimizations under certain key considerations. First, longer assay run times allow for farther diffusion of progeny virus from ground zero of a PCU infection. Second, less viscosity in the semi-solid overlay media increases flux of convection currents in the media, which also facilitates diffusion of progeny virus from ground zero of the PCU infection. As spillover was evident under conditions of 2% CMC and 19 hr assay runtime (**Supplementary Figure 2.2C-F, 2.3B-C**), we tested 8 hr, 10 hr, and 12 hr assay runtimes under 2% and 4% CMC. We found spillover to be mitigated under all tested conditions, however 8 hr and 10 hr were insufficient durations for PCUs to fully bloom and be counted as such, especially in slower growing strains like A/Texas/50/2012(H3N2) (**Supplementary Figure 2.4**). We also observed minor but inconsistent monolayer damage in the 2% CMC treatment groups, compared with no such damage in the 4% CMC groups. This damage was not observed during previous 19 hr assays, most likely because monolayer damage at the 12 hr time point had an additional 7 hr of recovery and resealing. Based on these findings, we settled on 4% CMC and 12 hr runtime for future assays involving A/California/07/2009(H1N1) and A/Texas/50/2012(H3N2) (**Supplementary Figure 2.5**).

**BIOHACKER ALERT**: Do not make CMC stock in microwave. Autoclave at 121°C for 30 min.

**2.3A**

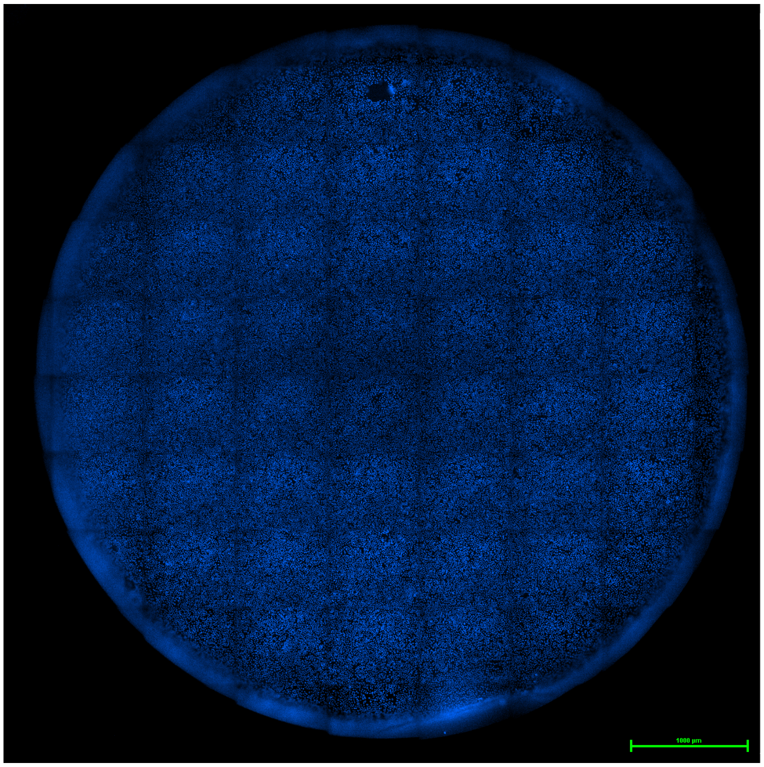


**2.3B**

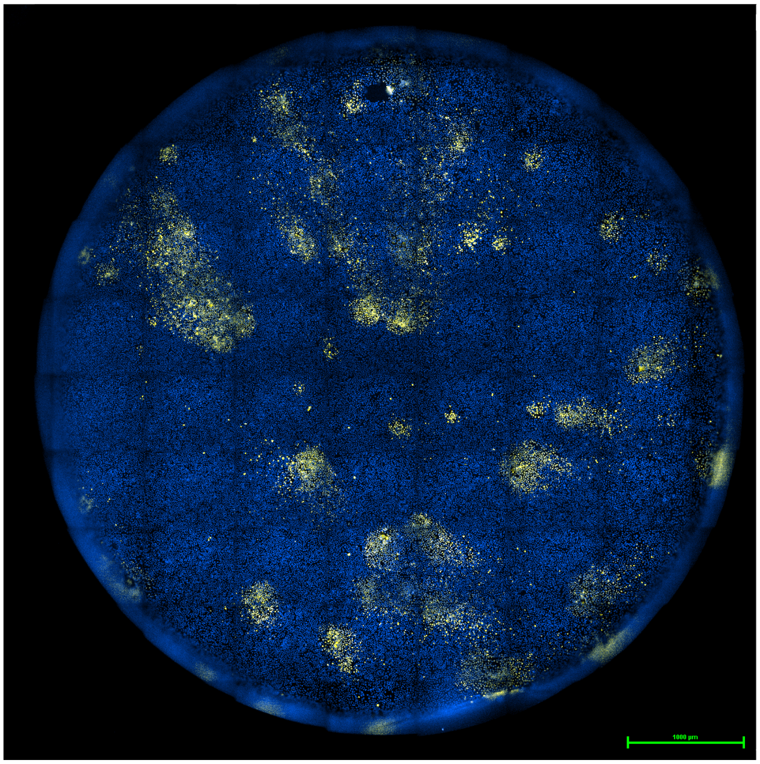

**2.3C**

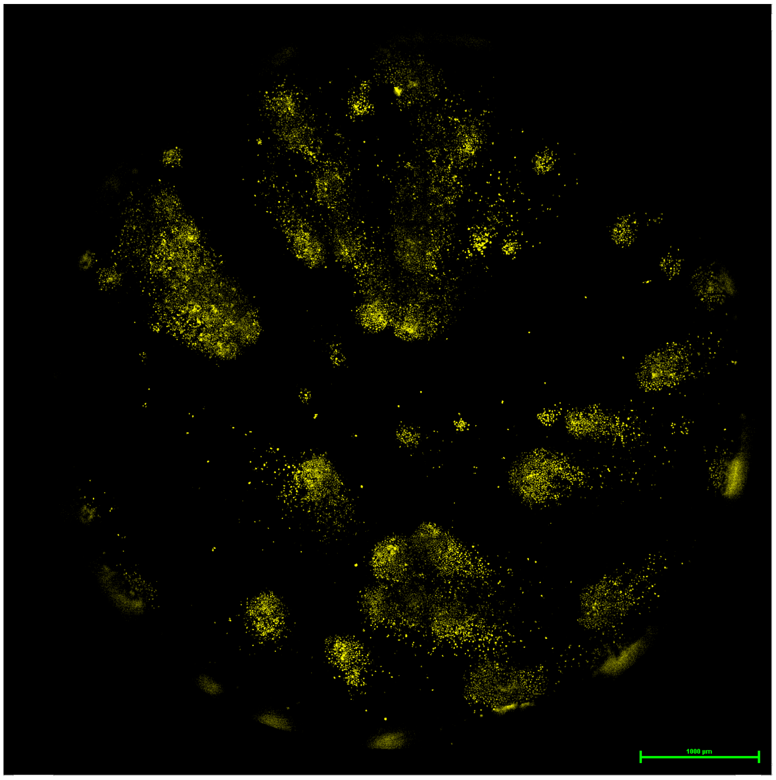

**Supplementary Figure S2.3.** (**A**) Barring aspirator pipette tip damage, a minimum cell seeding density (1E4 – 1.04E5 cells per well) is required to preserve monolayer integrity. However, PCU spillover and streaks still persist (**B-C**). Yellow/YFP – A/California/07/2009 nucleoprotein ; Blue/Hoechst – MDCK-London nucleus.

**2.4A**

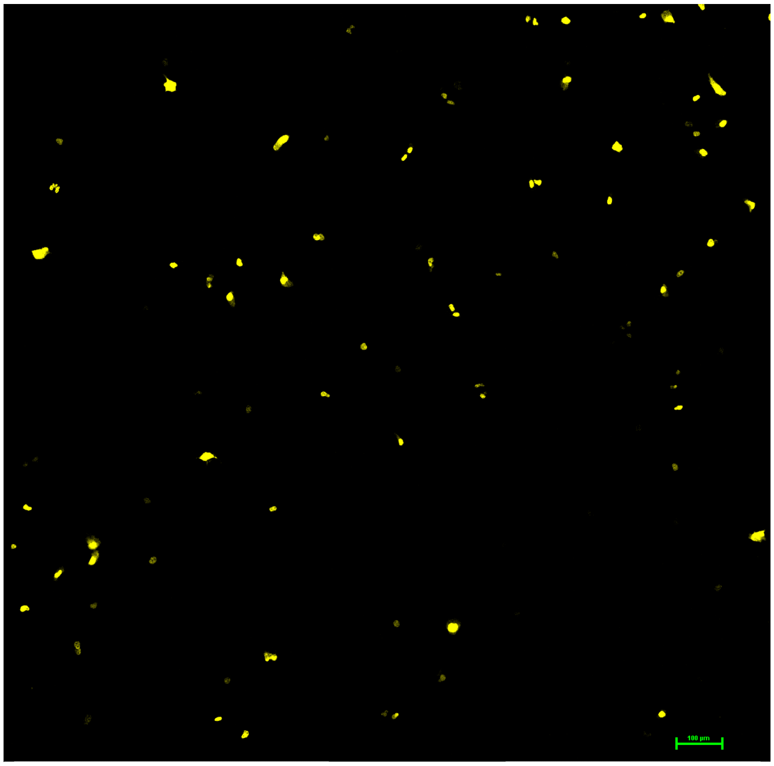

**2.4B**

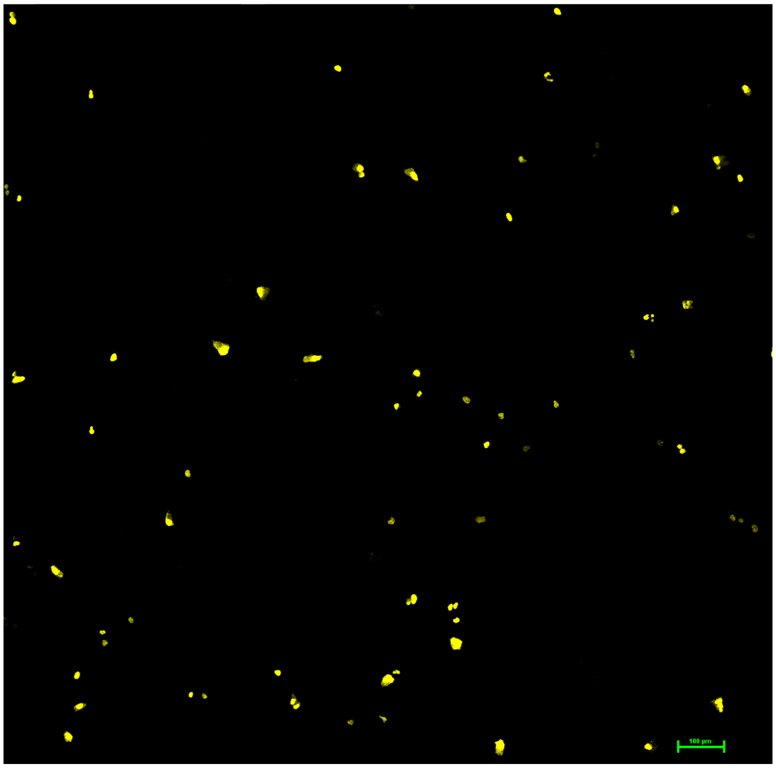


**2.4C**

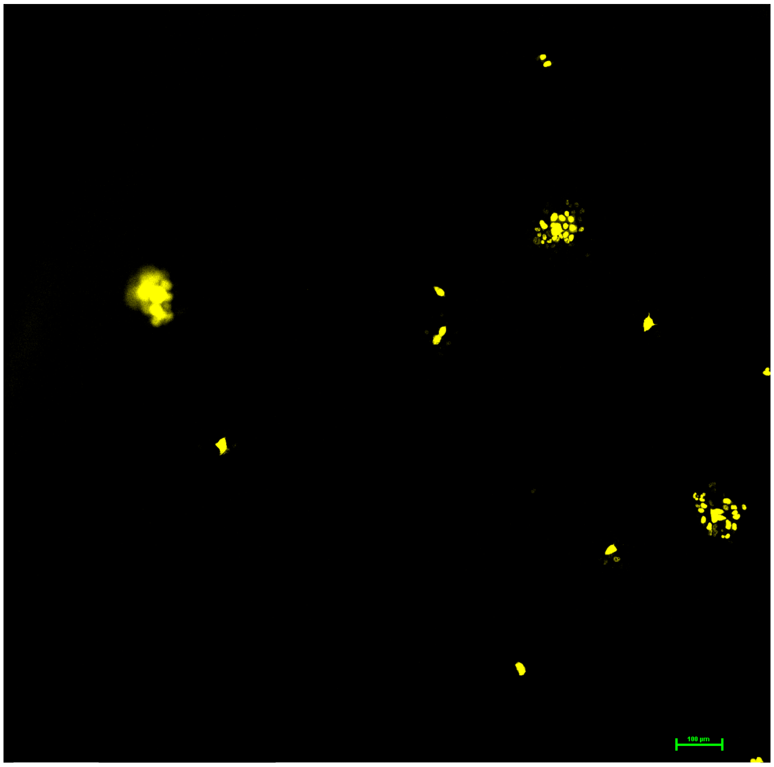

**2.4D**

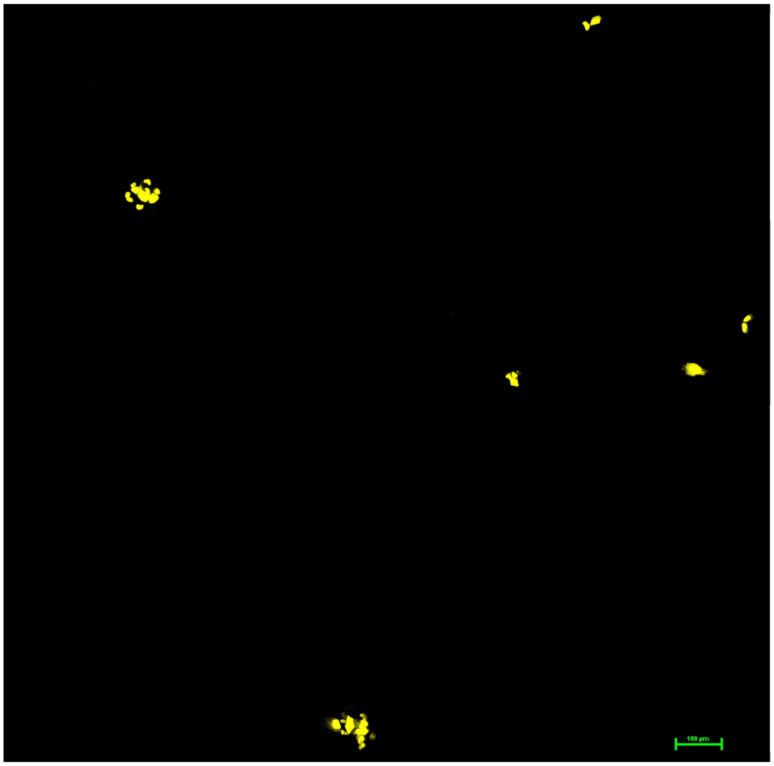

**Supplementary Figure S2.4.** Cluster-forming assay spillover mitigation; optimization of overlay concentration and assay runtime. (**A**) 8hpi at 2% CMC. (**B**) 8hpi at 4% CMC. (**C**) 12hpi at 2% CMC. (**D**) 12hpi at 4% CMC. Yellow(YFP) – A/California/07/2009 nucleoprotein.

**2.5A**

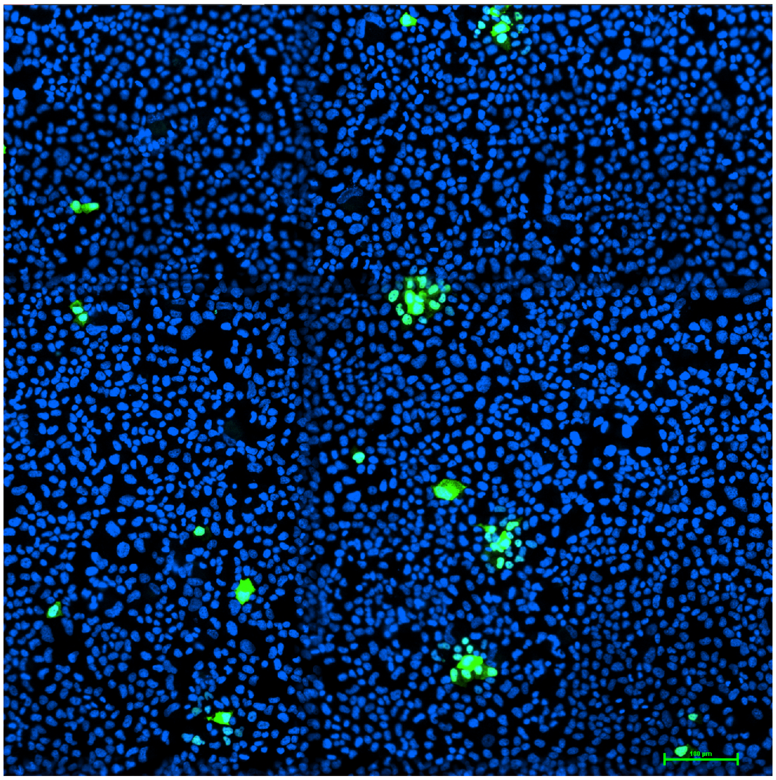

**2.5B**

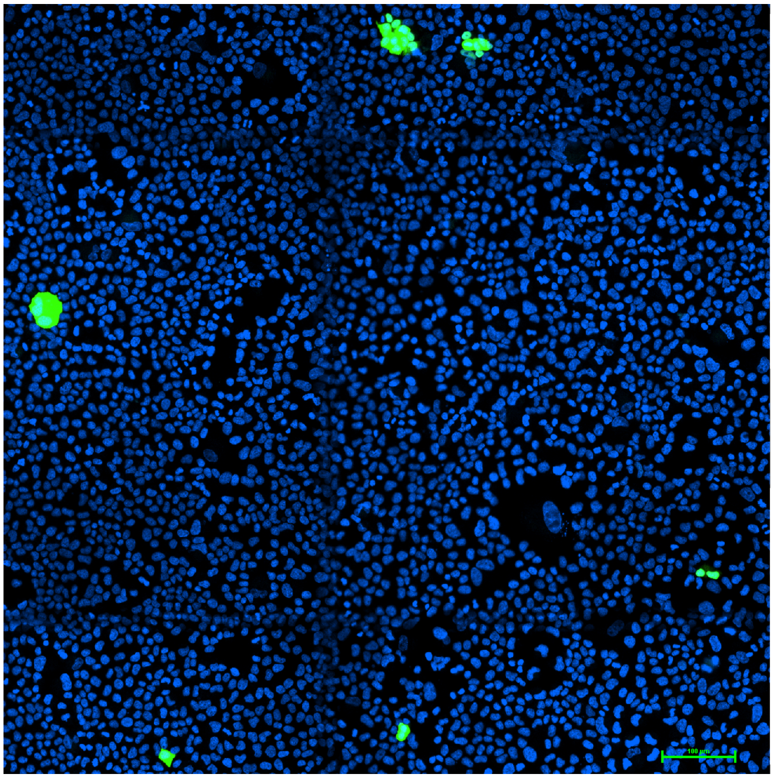

Supplementary Figure S2.5. Optimized cluster-forming assay; with damage-free monolayer and free of PCU spillover or streaks. Green/GFP – A/California/07/2009 nucleoprotein (**A**), A/Texas/50/2012 nucleoprotein (**B**) ; Blue/Hoechst – MDCK-London nucleus.

### *Cluster-counting Bioinformatics*

To count PCUs and NCUs, cluster-forming assay IF images (**Supplementary Figure 2.5**) were put through an automated image analysis pipeline we developed using MATLAB's image processing toolbox. Our guiding design principle was to cordon—or mask—nucleoprotein fluorescence signals (GFP) in the IF image as independent infection events, then overlay said mask with the host nuclei segmentation Hoechst signal to reveal the number of cells each infection event had spread to. We began stepwise assembly of masks around the GFP signals (**Supplementary Figure 2.6**) by binarizing IF images with the *imbinarize* function to make object detection possible, followed by the removal of small, noisy pixels with *bwareaopen*. Masks were sequentially dilated then filled with *imdilate* and *imfill* functions respectively to smoothen them out and ensure they did not contain holes. To finish the mask assembly, masks were eroded with *imerode* to undo the signal expansion done in the dilation step.

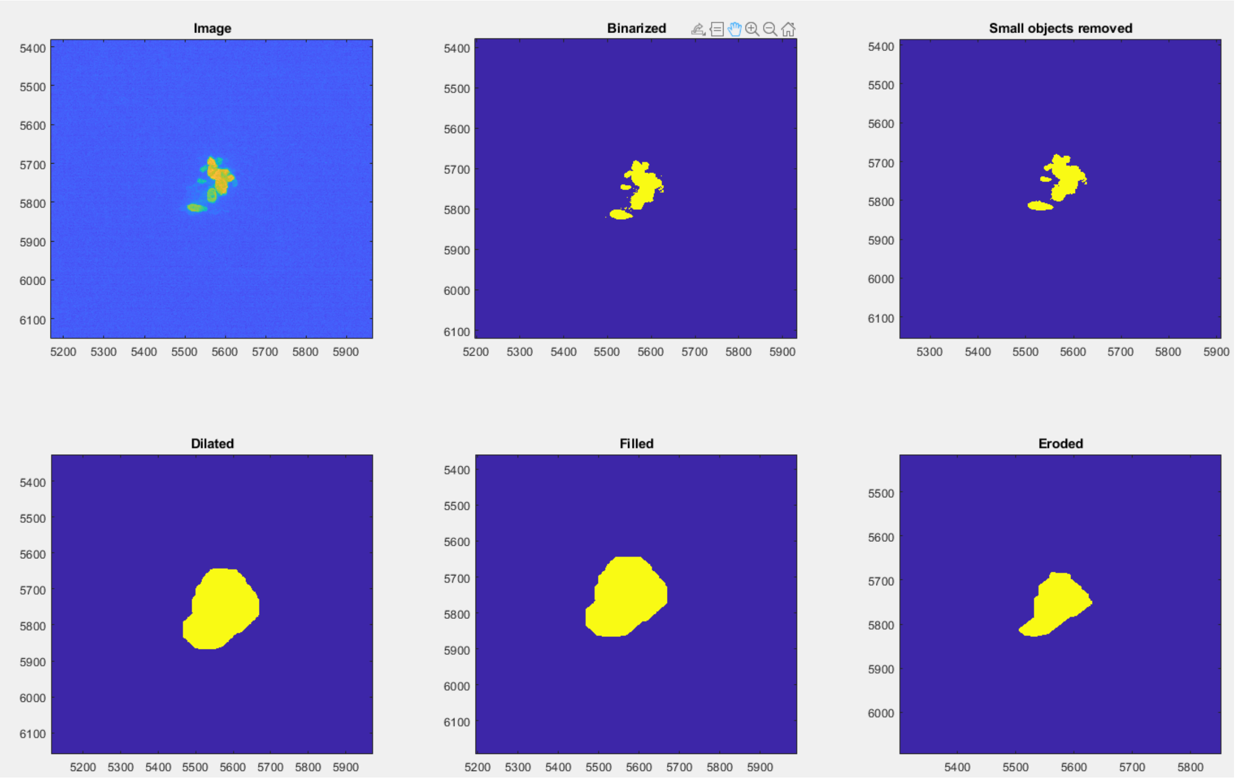


**Supplementary Figure S2.6.** Stepwise assembly of a mask around the nucleoprotein GFP signal in a productive clustering unit; starting from the initial cluster-forming assay IF image (**top-left**) down to final erosion (**bottom-right**).

At this juncture, the total number of masked objects in the binarized image included desired infection events, as well as undesired background noise from specs and autofluorescence. Undesired masks were filtered out by thresholding the min/max mask area and removing masks that did not contain any nuclei, leaving bona fide infection events—or clusters—that are counted and assigned a unique identity number (clusterID) (**Supplementary Figure 2.7**).

**2.7A**

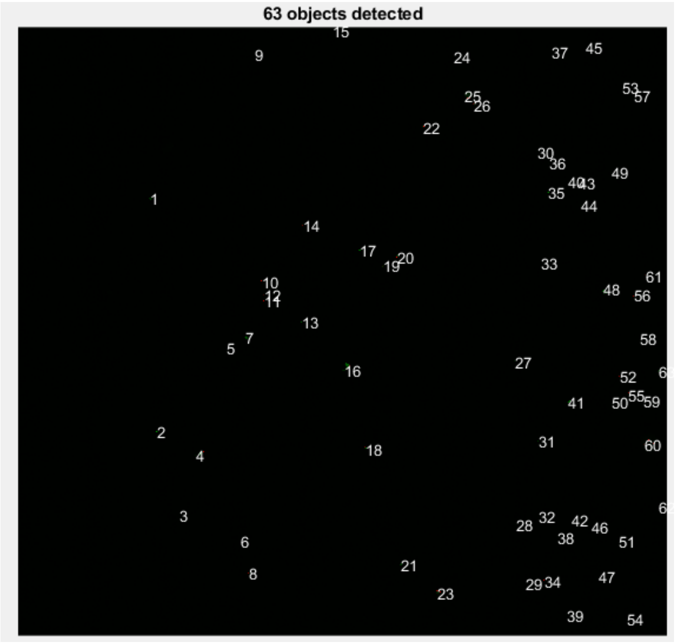

**2.7B**

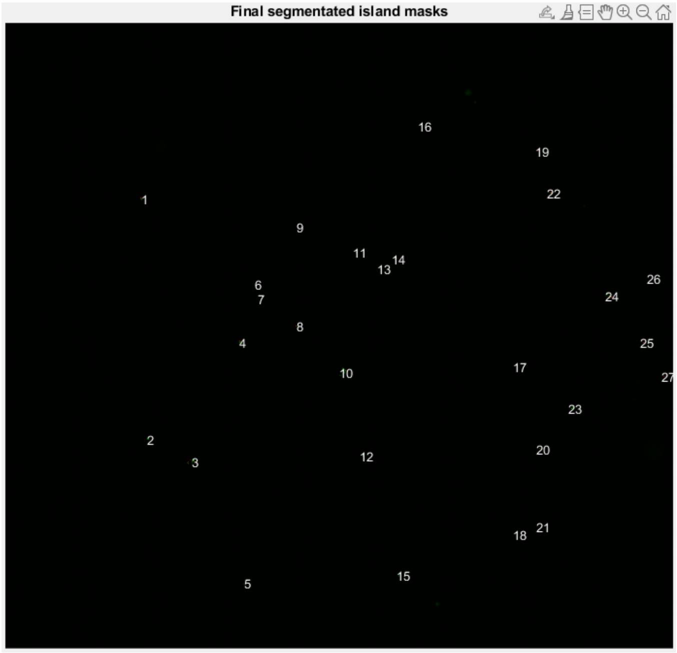


**Supplementary Figure S2.7.** Masked objects, before (**left**) and after (**right**) removal of background noise from specs and autofluorescence.

Next, segmentation was run on the Hoechst signal in host nuclei and overlaid with the masked clusters to demarcate and count the number of cells each infection had spread to (**Supplementary Figure 2.8**). Cluster data was sequentially exported to a spreadsheet then imported into the R Programming Language, where clusters were classified as PCU (ncell > 1) or NCU (ncell = 1), and after which NCU titer and NCU proportion were successively determined.

**2.8A**

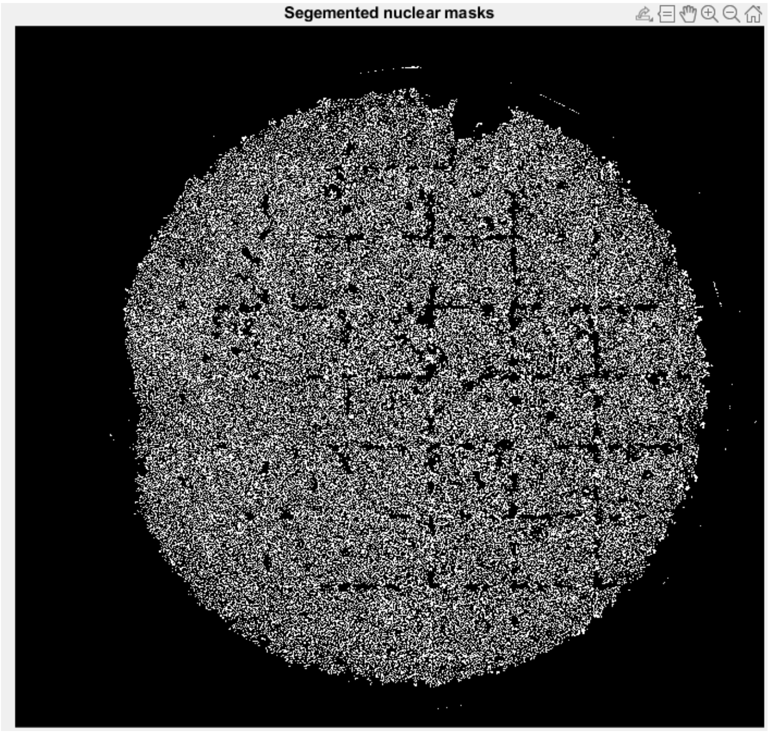

**2.8B**

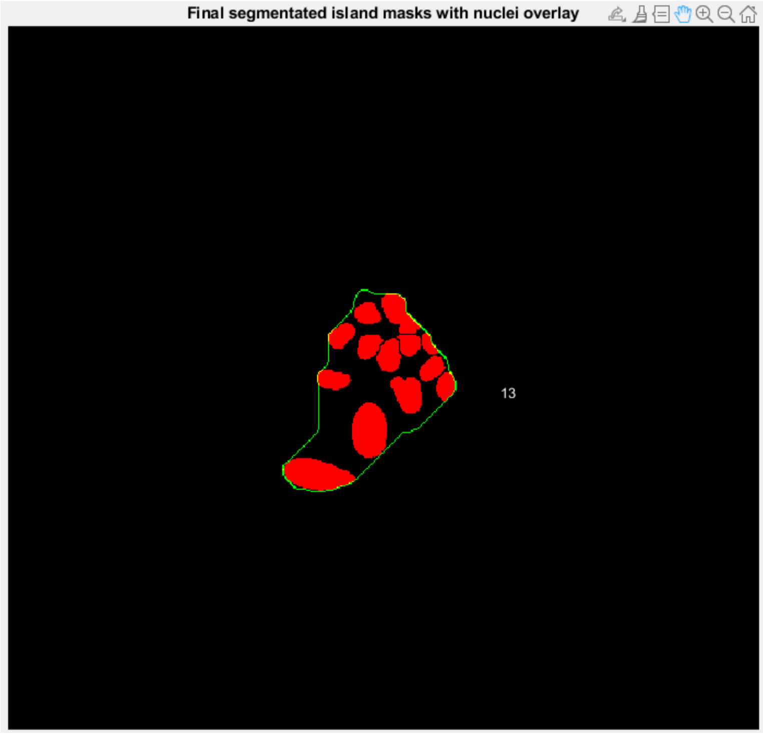


**Supplementary Figure S2.8.** Nuclear segmentation (**left**) and the segmentation-mask overlay (**right**) to size a clustering unit (PCU).
